# Simulation of FUS protein condensates with an adapted coarse-grained model

**DOI:** 10.1101/2020.10.10.334441

**Authors:** Zakarya Benayad, Sören von Bülow, Lukas S. Stelzl, Gerhard Hummer

## Abstract

Disordered proteins and nucleic acids can condense into droplets that resemble the membraneless organelles observed in living cells. MD simulations offer a unique tool to characterize the molecular interactions governing the formation of these biomolecular condensates, their physico-chemical properties, and the factors controlling their composition and size. However, biopolymer condensation depends sensitively on the balance between different energetic and entropic contributions. Here, we develop a general strategy to fine-tune the potential energy function for molecular dynamics simulations of biopolymer phase separation. We rebalance protein-protein interactions against solvation and entropic contributions to match the excess free energy of transferring proteins between dilute solution and condensate. We illustrate this formalism by simulating liquid droplet formation of the FUS low complexity domain (LCD) with a rebalanced MARTINI model. By scaling the strength of the nonbonded interactions in the coarse-grained MARTINI potential energy function, we map out a phase diagram in the plane of protein concentration and interaction strength. Above a critical scaling factor of *α_c_* ≈ 0.6, FUS LCD condensation is observed, where *α* = 1 and 0 correspond to full and repulsive interactions in the MARTINI model, respectively. For a scaling factor *α* = 0.65, we recover the experimental densities of the dilute and dense phases, and thus the excess protein transfer free energy into the droplet and the saturation concentration where FUS LCD condenses. In the region of phase separation, we simulate FUS LCD droplets of four different sizes in stable equilibrium with the dilute phase and slabs of condensed FUS LCD for tens of microseconds, and over one millisecond in aggregate. We determine surface tensions in the range of 0.01 to 0.4mN/m from the fluctuations of the droplet shape and from the capillary-wave-like broadening of the interface between the two phases. From the dynamics of the protein end-to-end distance, we estimate shear viscosities from 0.001 to 0.02Pas for the FUS LCD droplets with scaling factors *α* in the range of 0.625 to 0.75, where we observe liquid droplets. Significant hydration of the interior of the droplets keeps the proteins mobile and the droplets fluid.

## INTRODUCTION

Intracellular compartmentalization into organellar structures is crucial for the organization of the cellular biochemistry in time and space. The cell nucleus or mitochondria are examples of subcellular compartments bounded by a lipid membrane, whereas stress granules, processing bodies, nucleoli or Cajal bodies are membraneless.^1–6^ Disordered proteins and nucleic acids can cluster to form biomolecular condensates with liquid-like properties.^1,2,7,8^ Such condensates formed by liquid-liquid phase separation (LLPS) in vitro mimic the membraneless organelles in cells.^1,2^

Individually weak but multivalent interactions between intrinsically-disordered proteins (IDPs), in some cases amplified by condensing factors, are major drivers of LLPS.^4,8^ The biomolecular condensates produced by LLPS behave as liquid droplets immersed in dilute solution.^7^ Their liquid-like interior facilitates the rapid diffusion of reactants within the condensates and their exchange with the outside.^9^ Their dynamic nature also makes biomolecular condensates a promising template for novel biomimetic materials.^10,11^ The rational design of new materials will benefit from predictive models that relate the static and dynamic materials properties of biomolecular condensates to the protein and nucleic acid sequences^10^ and the molecular interactions they encode.

The RNA-binding protein Fused in Sarcoma (FUS) contains a low complexity domain (LCD) and is implicated in the formation of membraneless organelles.^12^ The FUS LCD has served as paradigm to understand biomolecular condensates.^13,14^ Enriched in the nucleus, FUS participates in transcription, DNA repair and RNA biogenesis.^15^ It has also been shown that disease-associated mutations lead to the liquid-to-solid transition of FUS droplets through the formation of fibrous amyloid-like assemblies.^9,16^

Theory and simulations^17–25^ contribute to the emerging understanding of LLPS. Molecular dynamics (MD) simulations at atomic resolution could in principle monitor molecular interactions with high accuracy. While high computational costs still preclude their widespread use for simulating LLPS, they promise considerable insights into structures and interactions of biomolecules in (sub-systems of) phase-separated condensates^26–29^ and how these give rise to materials properties.^30^ Soft-matter approaches^31^ provide access to the large length and long timescales relevant for phase separation, but may not resolve the detailed effects of protein chemistry and may not be transferable. Overcoming these two issues would require transferable models. ^20,21^ Coarse-grained simulations using a reduced representation^32^ enable direct simulations of phase behavior^21,24,33,34^ and can give a good description of effective molecular interactions, which is important for transferability. Dignon et al. ^21^ studied the phase separation of the low complexity domain of FUS, using a coarse-grained protein model with implicit solvent. They built the phase diagram in the temperature-concentration space and demonstrated the importance of specific molecular interactions between the IDPs.

The parameterization of coarse-grained simulation models for quantitative studies of phase separation is challenging. In the case of proteins in solution, intra-protein, proteinprotein, protein-solvent and solvent-solvent interactions have to be balanced with each other and with configurational and solvation entropy contributions associated with the degrees of freedom integrated out in the coarse-grained representation. Systematic errors in representing molecular interactions are often extensive, i.e., their contributions to the excess free energy difference per molecule between the phases tend to grow linearly with chain length. As a consequence, even small systematic imbalances become amplified. Incidentally, similar challenges^35–37^ are faced in simulations of protein self-assembly using highly optimized all-atom force fields. These classical force fields have been parameterized primarily using quantum mechanical data and, in this sense, are also coarse-grained. Indeed, tuning the helix-coil equilibrium of proteins ^35^ against experimental data turned out to be a critical improvement for simulations of de novo protein folding. ^38^ For disordered proteins, rebalancing of nonbonded and solvent interactions addressed the issue of compactness in disordered proteins.^36,37^

Here we adopt this strategy to develop and implement a general approach to fine-tune MD simulation models for studies of biopolymer phase separation. In a first step, we adjust the strength of the nonbonded interactions between the droplet-forming biopolymers to reproduce the experimental excess transfer free energy between the dilute and dense (droplet) phase. Reproducing the experimental densities of both the dilute and the dense phase gives us a first validation test, and an indication that the model may provide a reasonable description also of the molecular structures in the two phases. In a second step, we use the rebalanced model to calculate structural, thermodynamic and dynamic materials properties and to compare them to experiments where possible. We demonstrate this general procedure in simulations of the phase behavior of the disordered protein FUS LCD with the MARTINI model,^39^ a widely used coarse-grained force field.

We first map the phase diagram in the plane spanned by the protein concentration *c* and a parameter *α* introduced previously to scale the protein-protein Lennard-Jones (LJ) interaction energy. ^40,41^ Having identified the region where the FUS LCD undergoes spontaneous phase separation, we simulate stable FUS droplets at different protein-protein interaction strengths α. From these simulations, we construct the coexistence line in the *α-c* plane as the relationship between *α* and the protein concentrations c_dense_ and c_dilute_ in the droplet and in the coexisting dilute phase, respectively. Using the coexistence data, we determine the value of *α* for which the measured excess transfer free energy as well as the two densities are reproduced. We then quantify three biophysical properties: the hydration level of the droplets, their surface tension and their shear viscosity. The strong dependence of these structural, thermodynamic and dynamic properties on *α* demonstrates the importance of force-field rebalancing to describe biologically relevant behavior.

Our approach of rebalancing MD simulation force fields using experimental information on LLPS is not limited to a particular type of coarse-graining,^32^ but is generally applicable. Instead of or in addition to scaling protein-protein interactions, also other aspects of the potential energy surface can be adjusted. Force-field rebalancing to match phase boundaries and excess transfer free energies should improve the reliability in molecular simulations of biopolymer phase separation.

## SIMULATION METHODS

### Protein-protein interaction described by scaled MARTINI 2.2 forcefield

A variant of the MARTINI 2.2 force field^42^ was used for all MD simulations, in which we rescaled the protein-protein LJ interactions following Stark et al.^40^. They introduced a parameter *α* to scale the protein-protein LJ pair interaction well depth, *ϵ_α_* = *ϵ*_0_ + *α*(*ϵ*_original_ – *ϵ*_0_). A value of *α* = 0 corresponds to a repulsion-dominated interaction in the MARTINI model,^42^ *ϵ*_0_ = 2 kJ /mol, and a value of *α* =1 recovers the full interaction in the MARTINI force field, *ϵ*_1_ = *ϵ*_original_. Adopting a common rebalancing strategy for nonspecific protein-protein interactions, ^43,44^ Stark et al. ^40^ adjusted α to match experimentally determined protein-protein virial coefficients, which also gave promising results for carbohydrates.^45^ In the spirit of the MARTINI force field development,^39^ here we instead attempt to match the experimentally determined excess transfer free energy for a protein between the dilute phase and the dense phase of condensed droplets. In accordance with the method of Stark et al.^40^, interactions involving the beads P4, Qa, and Qd were not scaled, because they also describe the water and ion beads in the system.

### Simulations to map the phase boundaries in the *α-c* plane

In a first exploratory step, we qualitatively mapped the phase boundaries in the plane spanned by the protein concentration *c* and the interaction strength *α* using MD simulations. We initiated these simulations from homogeneous solutions of protein chains in a cubic box with periodic boundary conditions. All simulations were performed with the GROMACS^46,47^ v2018.6 software. In the simulations, we varied the protein concentration and the interaction strength *α*. The FUS LCD is 163 amino acids long. Its amino-acid sequence is listed in Table S1. We built an initial atomistic model of the FUS LCD in an extended conformation using the AmberTools Leap program,^48^ which we then coarse-grained using the `martinize.py` code.^42,49^ We built initial simulation systems by randomly inserting copies of the coarse-grained model into the simulation box and adding water beads. Sodium ions were added to ensure electroneutrality. We replaced 10% of the water beads with “anti-freeze” beads to prevent non-physical solvent freezing. In an initial equilibration, we used repulsion-dominated protein-protein interactions (*α* = 0) to disperse the proteins in the box. We performed a first MD simulation of 2 ns in the NVT ensemble, using the velocity scaling thermostat^50^ to establish a temperature of 300 K. A second equilibration step was conducted for 20 ns in the NPT ensemble at a pressure of 1 bar, maintained by an isotropic Berendsen barostat. ^51^ Production simulations were then started from the final conformation with dispersed FUS LCD. In the production simulations, *α* was set to the desired value between zero and one. At each *α* value, we performed simulations of 12 μs (see Table S2), using a time step of 30 fs. Temperature and pressure were maintained at 300 K and 1 bar, respectively, by a velocity scaling thermostat ^50^ and a Parrinello-Rahman barostat.^52^ The “New-RF” MD settings as described in ref^53^ were used with short-range LJ and electrostatics cutoffs of 1.1 nm. The trajectories were visually inspected using the VMD^54^ v1.9.3 software.

### Cluster formation

We examined the MD trajectories for possible phase separation by monitoring the formation of protein clusters. Protein clusters were identified on the basis of a pair-distance criterion. In a given simulation structure, two proteins were considered to be in the same cluster if any pair of beads was within a cutoff distance of 5 Å. We monitored the FUS LCD condensation process by following the number and size of protein clusters along the trajectory.

### Simulation of droplets

FUS LCD droplets were simulated over several microseconds by starting from a preformed droplet. Droplet formation kinetics directly from homogeneous solutions can be severely slowed down by periodic boundaries, which favor percolation in sufficiently dense systems. To avoid percolation, we reduced the protein concentration by increasing the box size and adding solvent. To simulate droplets in equilibrium with the dilute phase over several microseconds, we hence started from a droplet formed at *α* = 0.7 (see Table S2 for details). We then extended the trajectory at different α values. The production phase of the droplet simulations started after an equilibration at the target value of α lasting several microseconds. Table S3 lists the trajectory ranges used for surface tension calculations. For *α* = 0.625 near the critical value, we restricted the surface-tension calculation to the segments of the trajectory where the droplets were clearly discernible.

### Calculation of droplet density

We computed the radial density profile of MARTINI beads with respect to the droplet center of mass. We converted the number densities into mass densities using a molar mass of 17.168 kg/mol for the simulated FUS LCD. We used the clustering algorithm described above, defined the largest cluster as “droplet” and calculated the radial density profile with respect to its center of mass. The same method was used to compute radial water density profiles.

### Matching of the excess transfer free energy

The excess transfer free energy Δ*G*_trans_ to bring one protein chain from the dilute phase to the dense droplet phase was computed as follows:

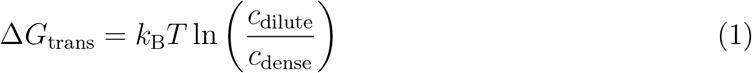

where *k*_B_ is Boltzmann’s constant, *T* is the absolute temperature, and *c*_dilute_ and *c*_dense_ are the concentrations (i.e., mass densities) in the dilute and dense phase, respectively. Here we neglected surface effects linked to the droplet curvature. The concentration of FUS in the droplet was calculated using the method described in the previous section. For the concentration of the dilute phase, we divided the number of proteins not part of the largest cluster by the box volume minus the volume of the droplet approximated as a sphere. The droplet radius was obtained from a sigmoidal fit of the protein-mass density profiles *c*(*r*),

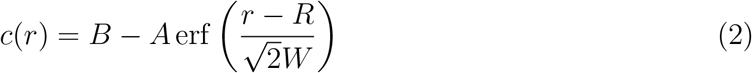

where *r* is the distance from the droplet center of mass; erf is the error function; and *A, B, R* and *W* are fitted parameters. Note that the derivative of this density profile is a Gaussian distribution with standard deviation *W*.^55^ For sufficiently large droplets with a defined density plateau at their center, *B* + *A* = *c*_dense_ and *B* – *A* = *c*_dilute_ correspond to the protein concentrations in the dense and dilute phases, respectively. From these limiting densities, we obtained excess transfer free energies for several *α* values that we compared to experimental results for the FUS LCD found in the literature.^13,14,27^

### Surface effects linked to the droplet curvature

We computed the excess transfer free energy for a flat surface using the slab simulation method.^56,57^ A dense phase forming a continuous slab under periodic boundary conditions was simulated in equilibrium with a dilute phase in an elongated box, with slab surfaces normal to the largest dimension (parallel to the z-axis). We identified the dense phase as the largest cluster, and considered all other proteins as part of the dilute phase. The concentration of the dilute phase was determined by dividing the number of proteins in the dilute phase by the volume remaining after subtraction of the slab volume, approximated as a rectangular parallelepiped of dimension *L_x_* × *L_y_* × 2*z*_0_. *L_x_* and *L_y_* are the box dimensions in the *x* and *y* directions respectively, and ±z_0_ are the midpoints of the mass density profiles centered on the origin and fitted to the sigmoidal function eq 2. The excess transfer free energy was computed according to eq 1.

### Determination of the surface tension

The surface tension was estimated using two methods, both based on the theory of Henderson and Lekner^58^ for the thermal fluctuations of the shape of a spherical droplet composed of an incompressible fluid. In this model, thermally activated capillary waves roughen the interface and result in fluctuations of the droplet shape. These thermal fluctuations are associated with small changes *δA* in the surface area of the droplet relative to the sphere. The potential energy *U* ≈ *γδA* of surface-shape fluctuations is assumed to be dominated by the surface tension *γ*. As derived in Supporting Information, the fluctuations in the shape of the droplet at lowest order in a spherical harmonics expansion give us two independent estimates of the surface tension,

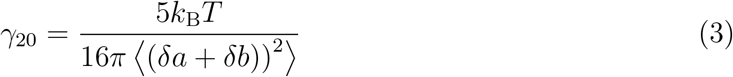

and

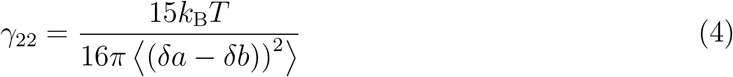

where *δa* = *a* – *R* and *δb* = *b* – *R* are the differences in lengths of any pair of principal axes of a general ellipsoid describing the instantaneous shape of the droplet with respect to the average radius of the droplet, *R*. The average 〈·〉 is thus both over the droplet shapes along the MD trajectory and over the three distinct combinations of principal axes. We estimated the three axis lengths *a, b* and *c* from a principal component analysis (PCA) of the mass distribution, as described in Supporting Information.

We obtained a third estimate of the surface tension from the width of the interface between the droplet and the dilute phase, again relying on the theory of Henderson and Lekner^58^. Capillary-wave theory predicts that the squared interface thickness *W*^2^ of an incompressible droplet grows as the logarithm of the droplet radius, with a prefactor proportional to the reciprocal of the surface tension,

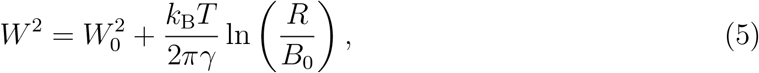

where *W* is the interface width, *W*_0_ is an intrinsic width, *R* is the droplet radius, and *B*_0_ is a short wavelength cutoff. Mittal and Hummer^55^ used this relation to extract an interfacial tension for the water-vapor interface around a hydrophobic solute. ^55^ Following their procedure, we simulated droplets of different diameters by varying the number of proteins. We extracted the interfacial width *W* of the droplets from fits of eq 2 to the radial concentration profiles. The surface tension was then determined from the slope of a linear fit to *W*^2^ as a function of ln *R* according to eq 5. Standard errors of *W*^2^ and ln *R* were estimated from block averaging.

### Estimation of the shear viscosity

To estimate the shear viscosity *η*, we computed the normalized autocorrelation function of the end-to-end distance of the protein chains. We estimated the autocorrelation time *τ* of the end-to-end distance as the amplitude-weighted sum of the two relaxation times in a bi-exponential fit to the autocorrelation function averaged over all proteins in the system. We approximated the effective diffusion coefficient *D* of the end-to-end motion as the ratio of the variance of the end-to-end distance *r* and the autocorrelation time *τ*,

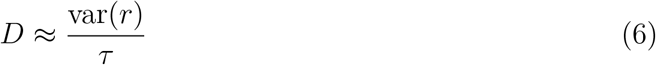

This expression becomes exact in the harmonic limit^59^ and accounts for possible effects of *α* on the compactness^24^ of the FUS LCD chains within the dense phase (Figure S1). For reference, we performed the same analysis for isolated FUS LCD chains in aqueous solution, giving us diffusion coefficients *D*_0_. We then assumed that the effective diffusion coefficients for the end-to-end distance relaxation scale as 1/*η*. Accordingly, we estimated the shear viscosity as

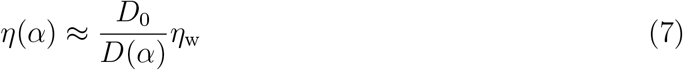

As reference, we used the effective end-to-end diffusion coefficient *D*_0_ of an isolated FUS LCD chain with *α* = 0.6, where we expect the contributions of attractive non-bonded interactions to the friction to be small,^60^ such that solvent contributions dominate. For the viscosity of MARTINI water, we used *η*_w_ = 1.0 × 10^−3^ Pas.^61^ With proteins frequently transferring between the droplet and the dilute phase, our analysis does not distinguish if a particular protein is inside or outside the droplet. However, only at values of *α* close to 0.6, where the droplet dissolves, contributions from proteins in the dilute phase become significant. In the most relevant regime of *α* ≥ 0.65, nearly all proteins are within the droplet and eq 7 thus reports on the viscosity within the droplet.

## RESULTS

### Phase separation of FUS LCD

We simulated ensembles of FUS proteins at different protein numbers, box sizes and interaction scaling factors *α*, with homogeneous solutions or preformed droplets as starting configurations, as detailed in Table S2. Above a critical interaction strength, *α* > 0.6, we observed phase separation at all protein concentrations simulated, 11 to 300mg/mL (Figure 1). In simulations starting from configurations with dispersed proteins at low average protein concentration, phase separation led to the formation of a roughly spherical droplet consistent with experimental studies of FUS.^9,62,63^ For *α* > 0.6, FUS LCD also formed condensates in MD simulations at high concentration of initially dispersed proteins. However, the topology of the resulting dense phase was inverted. For *α* > 0.6 and concentrations *c* > 100mg/mL, continuous protein condensates formed that percolated across the periodic boundaries of the simulation box (see inset at top right of Figure 1).

**Figure 1:**
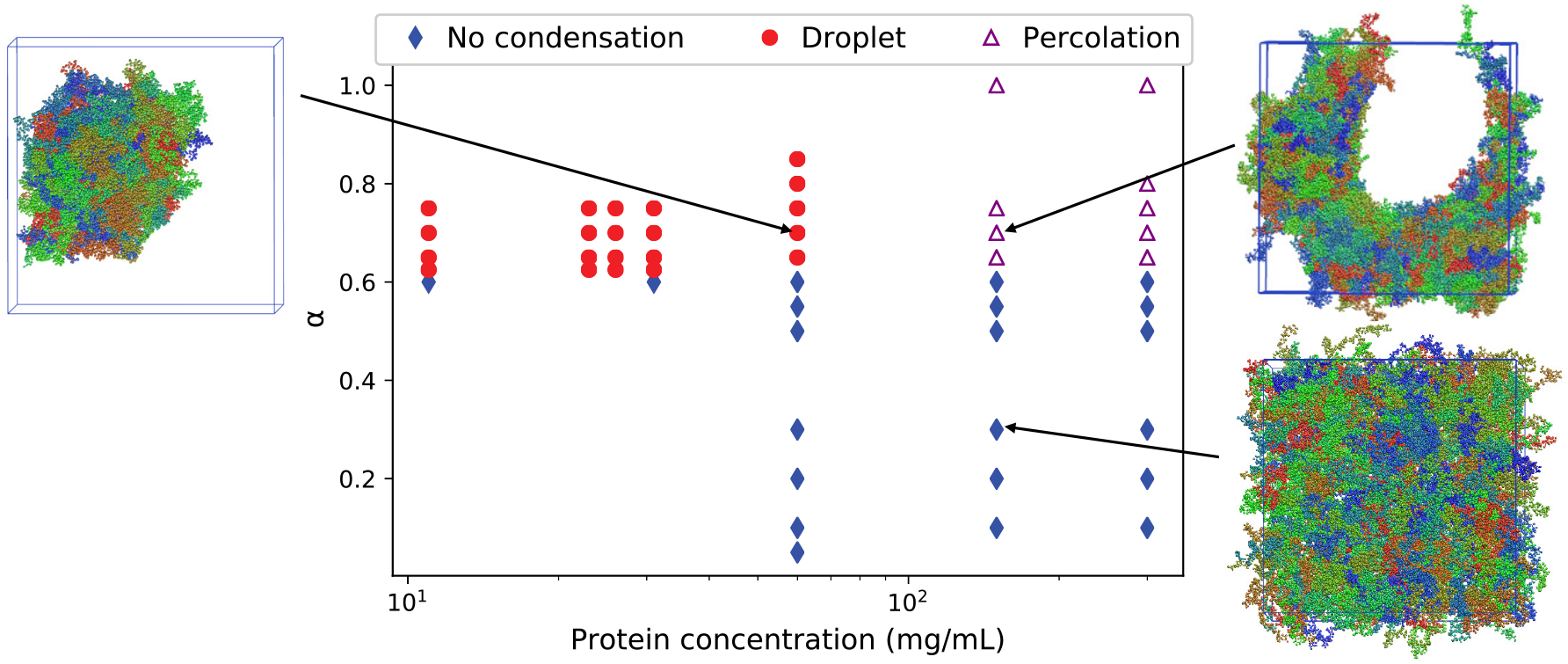
Phase diagram of FUS LCD. Symbols indicate the morphologies observed in MD simulations at different values of the protein concentration and the scaling parameter α (blue filled diamonds: dispersed proteins without condensation; red filled circles: droplets; open triangles: dense protein aggregates percolated across simulation box boundaries in an inverted topology). Insets show representative structures, connected by arrows to the respective point in the α-c plane. Shown are the FUS LCD chains in different colors, with water and ions omitted for clarity. The simulation results correspond to the “homogeneous” starting configuration in Table S2 for concentrations of 150mg/mL or higher and for 60mg/mL at *α* ≤ 0.7, and to the “preformed droplet” simulations for the rest (see Table S2).

### Cluster formation

We monitored the formation of FUS LCD clusters to characterize the process of phase separation and to identify droplets. As we increased the interaction scaling factor *α* beyond 0.6 at fixed protein concentration and temperature, we found the system to phase separate through the formation of clusters. Supporting Movie S1 shows the formation of a droplet starting from a homogeneous solution of 134 proteins at *α* = 0.7. The proteins first condense into small clusters, which then merge to form a droplet. For 0.6 < *α* ≤ 0.7, FUS LCD condensation leads to the coexistence of a dense phase and a dilute phase.

### Critical interaction strength

To quantify the critical interaction strength, we measured the time-averaged number and size of clusters at different values of *α* (Figure 2). The cluster analysis was performed on simulations starting from a homogeneous solution of 134 proteins in a 40 × 40 × 40 nm^3^ box at different *α* values (Table S2, Figure S2). At low *α* values (*α* < 0.6), the systems did not undergo phase separation. We found that the proteins remained dispersed, forming only small clusters that each contained only a small fraction of the proteins (Figure S2B). As *α* increases beyond 0.6, the number of clusters drops while the average cluster size grows sharply (Figure S2A,B). Phase separation led to the formation of one big cluster containing most of the proteins (> 95% for *α* ≥ 0.65) in coexistence with a dilute phase. The variations of these quantities with *α* indicate *α_c_* ≈ 0.6 as the critical interaction strength for our FUS LCD model at ambient temperature and pressure, above which phase separation occurs.

**Figure 2:**
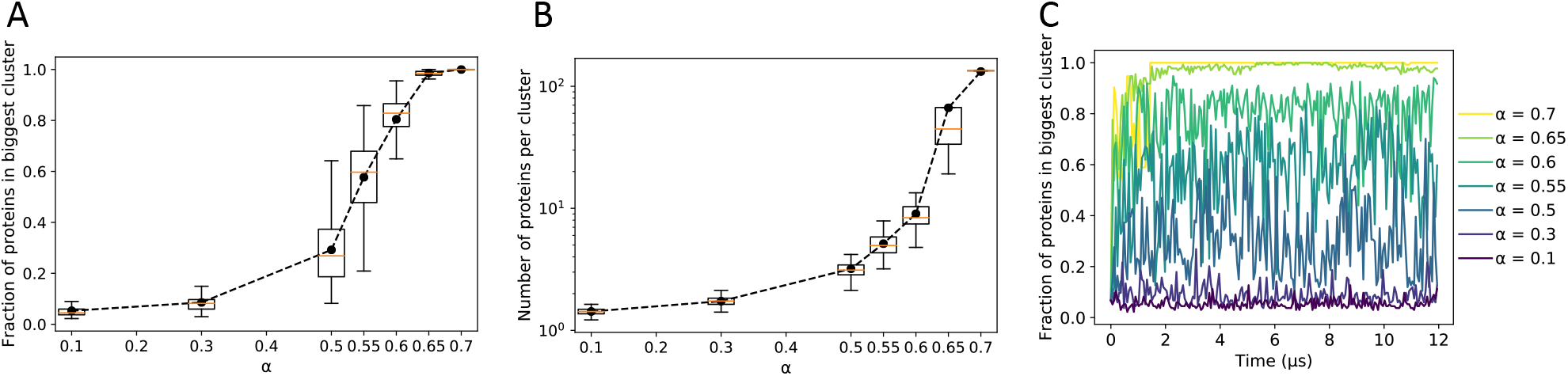
Cluster analysis of protein condensation as a function of the scaling parameter α. (A) Box-whisker plot of the fraction of proteins contained in the biggest cluster in a simulation (mean: filled circle; median: horizontal line; box: interquartile range; error bars: range). (B) Box-whisker plot of the number of proteins per cluster. (C) Fraction of proteins in the largest cluster as a function of time in MD simulations at different values of *α*, as indicated in the legend.

### Reversibility of phase separation

We confirmed that droplet formation is reversible. We initiated simulations with a droplet that had been formed at *α* = 0.7. After decreasing *α* to 0.55 and 0.5, we found the droplets to dissolve (Figure S3). This dissolution of preformed droplets at *α* < 0.6 is consistent with the phase behavior seen in simulations started with dispersed proteins. Droplet formation is thus reversible on the MD timescale.

### Density profiles

For *α* ≥ 0.65, phase separation led to the formation of a single droplet that remained stable over times >10 μs. We computed the protein density profiles inside the droplets at different *α* values (Figure 3A and Figure S4) from which we extracted the dependence of the dense-phase concentration on *α* (Figures 4A and 4C). We found that the protein density inside the droplets for a given *α* value is independent of the overall protein concentration in the system, consistent with the condensate being a distinct phase. The relevant quantities are hence *α* and the number of proteins *N*, which define the droplet density and size, respectively. The dense-phase concentrations for 0.65 ≤ *α* ≤ 0.7 are in the range of 300 to 400 mg/mL (Figure 4C), in line with dense-phase concentrations of IDPs reported from experiments.^27^ For *α* ≥ 0.8, we observed density inhomogeneities inside the “droplets” (Figure S4) that persisted on the simulation timescale. These irregular structures indicate that the strong protein-protein interactions for *α* ≥ 0. 8 slowed down the relaxation process beyond the timescale of our MD simulations. In the following, we focus our attention on the *α* region where phase separation leads to the formation of stable droplets with liquidlike properties.

**Figure 3:**
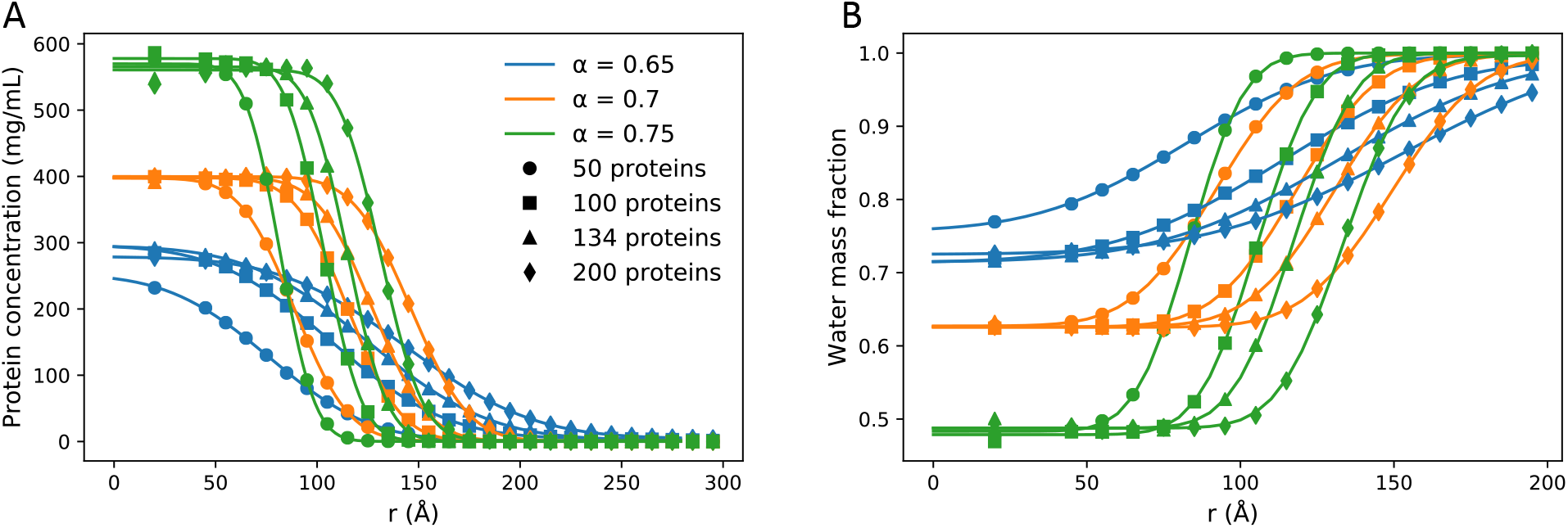
Radial protein and water concentration profiles in FUS LCD droplets. (A) Protein mass density as function of the radial distance *r* from the droplet center. (B) Relative mass fraction of water as function of *r*. Radial density profiles from MD simulations (symbols) are shown for different values of *α* values and different droplet sizes, as indicated. Lines are fits to the error function profile eq 2.

**Figure 4:**
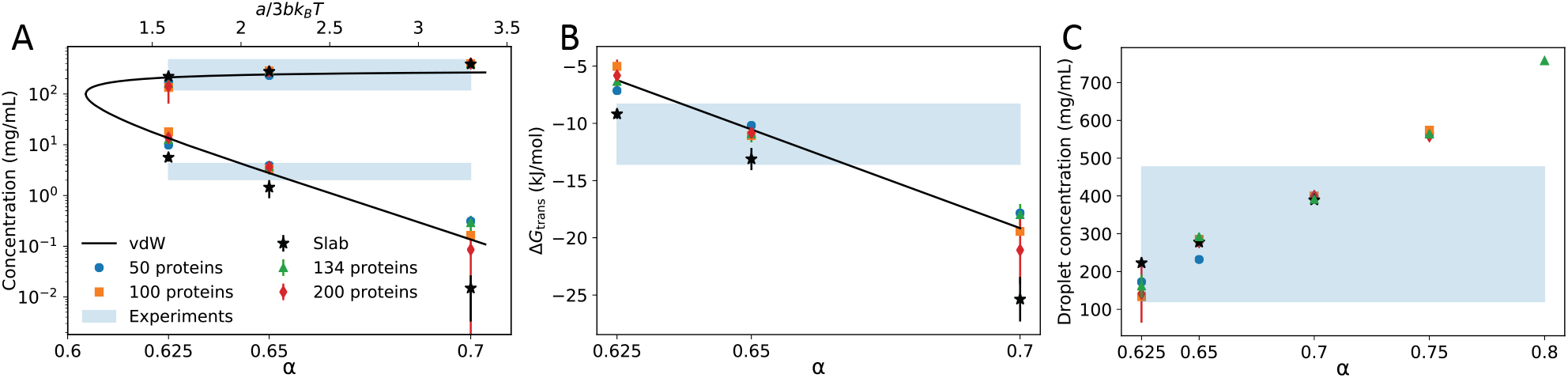
Phase diagram and excess transfer free energy for FUS LCD. (A) Concentration of the coexisting dense and dilute phases from MD simulations (symbols) for different values of *α* and droplet sizes, as indicated. The star markers show the densities obtained from the slab simulation method. ^56,57^ Bars for the largest droplets and the slab systems indicate standard errors. The black line shows the coexistence curve of van-der-Waals mean-field theory with *a*/(3*bk*_B_*T*) a linear function of *α*, as indicated on the top axis. The suitably adjusted critical density 1/(3*b*) corresponds to a critical protein concentration of 100mg/mL. Blue shading indicates the range of experimental values reported for the saturation concentration of FUS LCD^13,14^ and for the dense-phase concentration.^13,27^ (B) Excess transfer free energy of FUS LCD between the dilute phase and the dense phase (i.e., the droplet) calculated from eq 1. The black line is a fit to a linear function in *α*. Blue shading indicates the range of excess transfer free energies calculated for the experimental densities, as indicated by shading in (A). (C) Concentration of the dense FUS LCD phase as a function of the energy scaling parameter *a*. The blue area indicates the range of FUS LCD concentrations in condensates reported from experiment.^13,14^

### Excess transfer free energy between dilute and dense phases of FUS LCD condensate

The MARTINI force field^39^ has been parameterized extensively on the basis of excess transfer free energies (e.g., the water/oil partition coefficient). Adopting this general parameterization approach, we calculated the excess transfer free energy for FUS LCDs between coexisting dilute and dense phases. We used the cluster algorithm and the droplet density profiles to compute the protein concentrations of the two phases at coexistence. From the logarithm of their ratio, according to eq 1, we then obtained the excess transfer free energy.

Figure 4 shows the resulting densities and excess transfer free energies for different values of *α*, as obtained from simulations with different numbers of proteins and, thus, with different droplet sizes. We find that the density of the dilute phase changes by several orders of magnitude in the range of 0.625 ≤ *α* ≤ 0.7. By contrast, the protein concentration in the dense phase varies only by a factor of three in this α range.

For a value of *α* = 0.65, the densities of both the dense and the dilute phase, and so the excess transfer free energy, agree well with experimental values found in the literature.^13,14,27^ We therefore suggest *α* = 0.65 as a suitable scaling factor to describe the thermodynamics of FUS LCD phase separation at ambient conditions.

### Phase diagram and effective van-der-Waals fluid

The coexistence behavior in the *α-c* plane appears to be captured by van-der-Waals mean-field theory, ^64^ at least at a semi-quantitative level. As shown in Figure 4, a linear relation between the van-der-Waals interaction parameter *a* and *α* and a simple scaling of the protein concentration *c* bring the observed densities at coexistence into good correspondence with the coexistence curve of van-der-Waals theory in the plane spanned by the interaction strength *a* and density *ρ*. Here we fixed the temperature *T* and the volume parameter *b* in the van-der-Waals equation of state, *p* = *k*_B_*T*/(*v* – *b*) – *a/v*^2^, with *p* the pressure and *v* = 1/*ρ* the volume per particle. This correspondence is consistent with the estimate *α_c_* ≈ 0.6 of the critical value of the scaling parameter, below which no condensation occurs. A quantitative analysis of this correspondence would require careful finite-size scaling, which is beyond the scope of this work.

### Droplet composition

The presence of explicit solvent in the MARTINI model enabled us to characterize the hydration level of the droplet interior for different *α* values (Figures 3B and 5). For *α* = 0.65, water accounts for 70% of the droplet mass. At increasing *α*, the hydration level decreases. In effect, tightening the protein-protein interactions within the droplet squeezes out the water. Supporting Movie S2 visualizes the water inside the droplet for *α* = 0.7.

**Figure 5:**
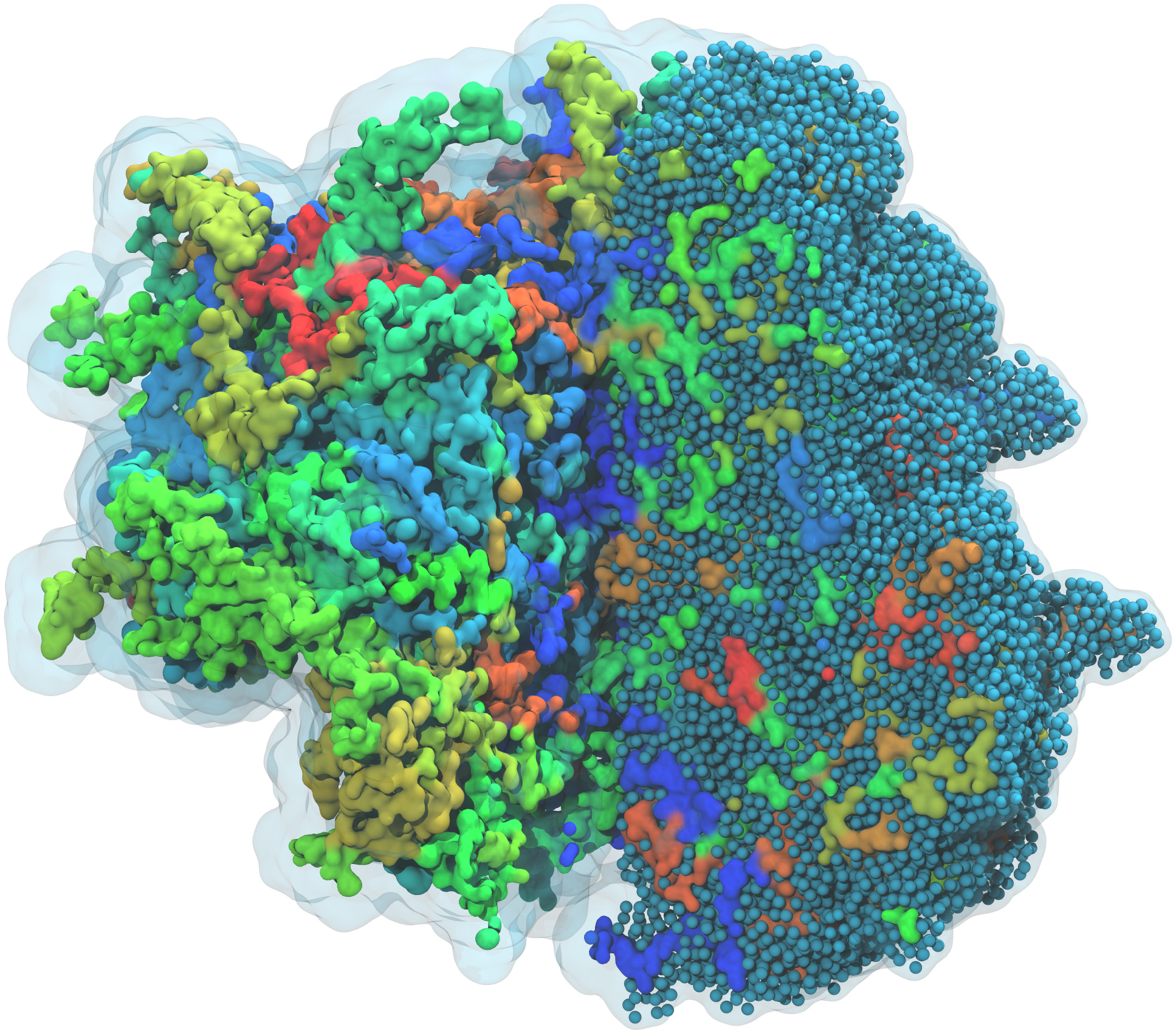
Hydration of the interior of FUS LCD droplet for *α* = 0.7 with *N* = 134 proteins. In this MD simulation snapshot, FUS LCD proteins in the left half of the droplet are shown in colored surface representation, with a transparent surface indicating water. To indicate the extent of hydration, water beads within the FUS LCD droplet, or at its surface, are shown as blue spheres in the right half of the droplet. The front right wedge has been removed to provide a clearer view of the droplet interior.

### Surface tension of FUS LCD droplets from capillary wave theory

We found that the surface tension of the FUS LCD droplets calculated from the interfacial width increases with *α* (Figure 6B). As shown in Figure 6A, the squared width of the droplet interface grows linearly with the logarithm of the droplet radius, as predicted by capillary wave theory, eq 5. In Figure S5, we show the error-function fits to the interfacial density profiles, from which the droplet interfacial widths and radii were extracted. Figure 6B shows the surface tension values calculated from the interfacial widths according to eq 5. At *α* = 0.625, where the critical *α* is approached, the surface tension is about 25 times lower than at *α* = 0.75.

**Figure 6:**
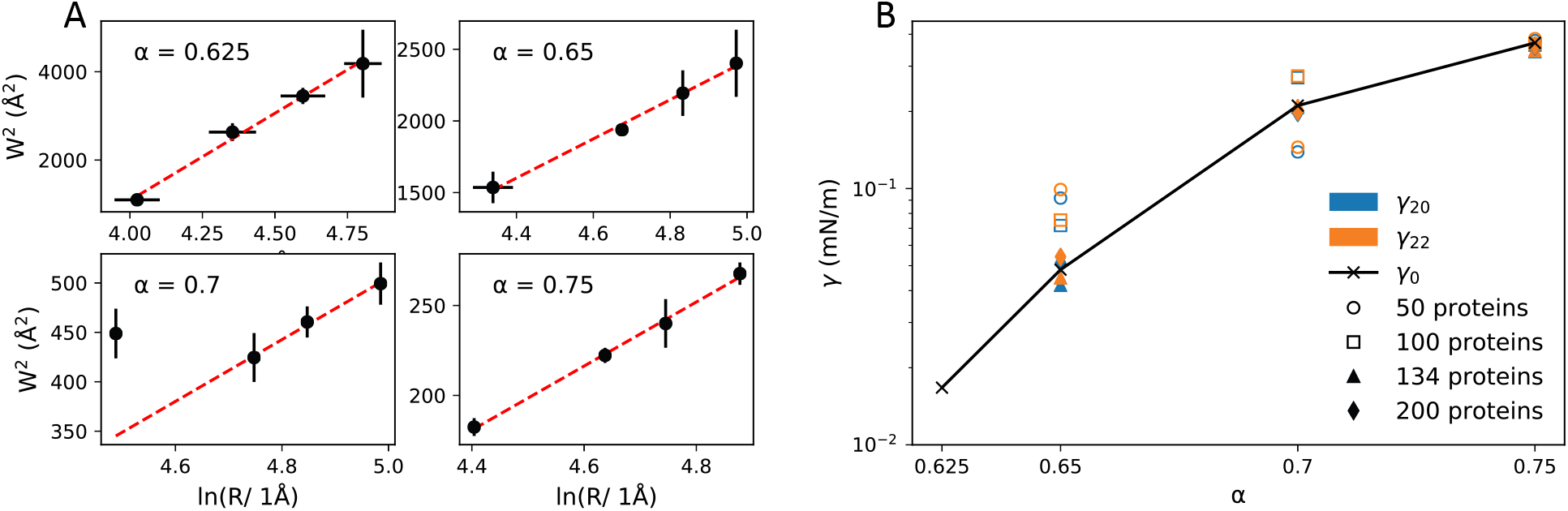
Surface tension of FUS LCD droplets. (A) Squared width of droplet interface as function of the logarithm of the droplet radius (symbols). Dashed lines are linear fits according to eq 5. For *α* = 0.7, the smallest droplet was considered an outlier and was thus not included in the fit. Standard errors, as indicated by error bars, were estimated by block averaging. Values of *α* are indicated. (B) Surface tension as function of *α* from eqs 3 (*γ*_20_) and 4 (*γ*_22_; filled and open symbols; droplet size is indicated), and from eq 5 (*γ*_0_; crosses connected by lines).

The surface tensions calculated from droplet shape fluctuations and from the interfacial width are consistent. Droplet shape fluctuations are illustrated in Figure 7C-F. For 0.65 ≤ *α* ≤ 0.70 and droplets with >100 proteins, where shape fluctuations could be resolved and adequately sampled, we obtained excellent agreement with the estimates from the interfacial width. For *α* = 0.625, poorly defined droplet shapes did not allow us to extract reliable ellipsoid axes. For *α* = 0.65, where the FUS LCD excess transfer free energy is matched, the surface tension is consistently about *γ* ≈ 0.05 mN/m.

**Figure 7:**
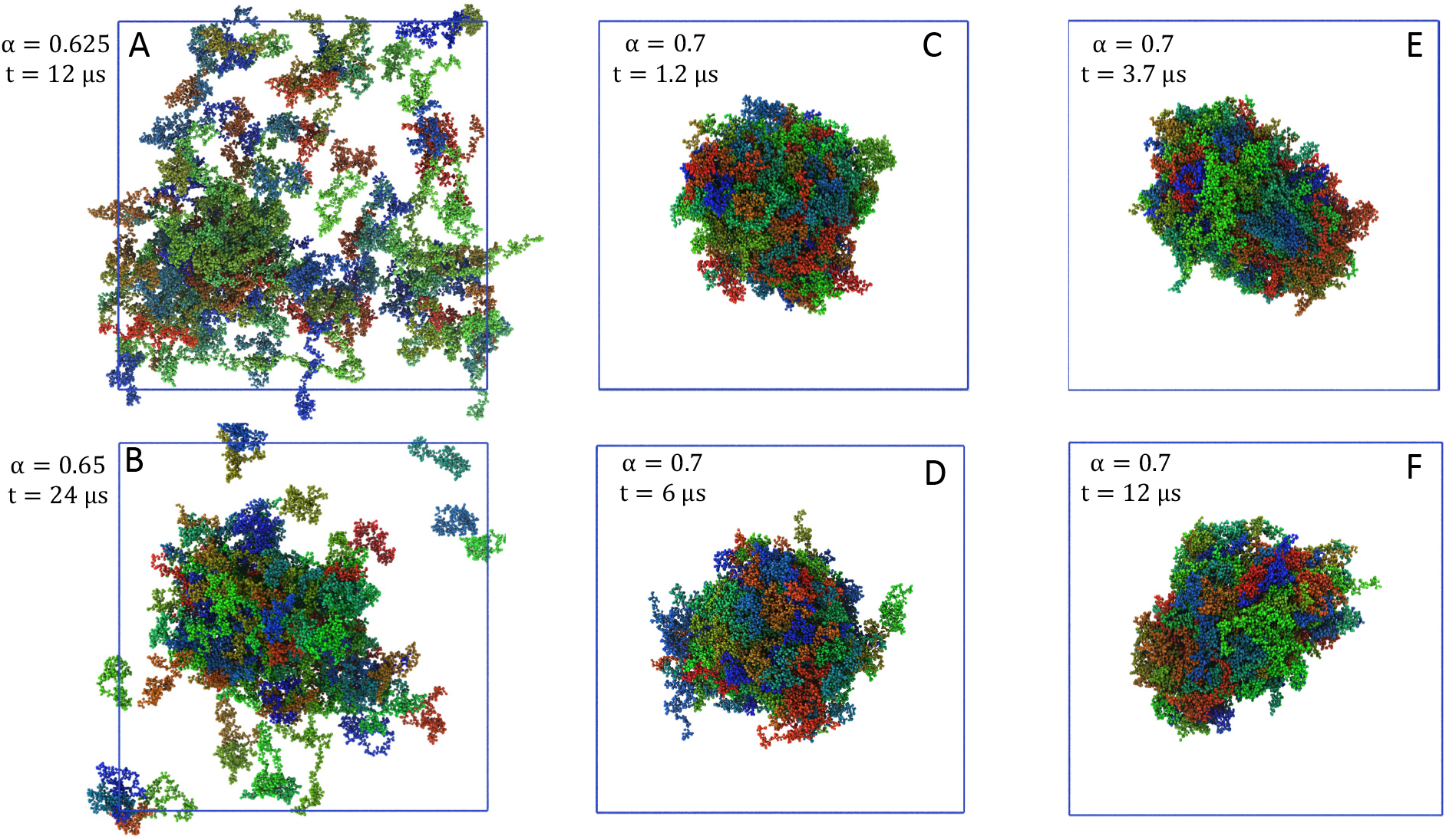
Snapshots of the MD simulation systems at different values of *α* and different time points. (A) *α* = 0.625. (B) *α* = 0.65. (C-F) *α* = 0.7 with time points indicated. The blue squares indicate the simulation box size.

As a further test of capillary wave theory, we compare in Figures S6 and S7 the distributions of the squared amplitudes of the ellipsoidal droplet shape fluctuations to the predicted exponential distributions: we have good correspondence, in line with the agreement in the surface tension values from droplet shapes and interface widths.

### Droplet shear viscosity from FUS end-to-end distance relaxation

We estimated the shear viscosity of the protein droplets from the standard deviation of the protein end-to-end distance and its relaxation time according to eq 7. We observed that the relaxation time *τ* depends exponentially on *α*, increasing by about a factor of ten between *α* = 0.6 (where droplets start to form) and *α* = 0.75 (Figure 8A). By contrast, the relaxation times of isolated FUS LCD chains free in solution are nearly independent of α. From the ratio of the standard deviation and the relaxation time (Figures S8 and S9), we obtained the end-to-end diffusion coefficient using eq 6. Then, we used eq 7 to calculate the viscosity, assuming that the diffusion coefficient scales as 1/*η*, with a reference effective end-to-end diffusion coefficient *D*_0_ that we had estimated from MD simulations of a free FUS chain at *α* = 0.6 (Figure S10). The calculated viscosities are shown in Figure 8B. We find that *η* increases exponentially with *α* from about 0.001 Pas to 0.02Pas. For *α* = 0.65, where our rebalanced model matches the experimental excess transfer free energy, we estimate a shear viscosity for FUS LCD droplets in the range of 0.002 to 0.004Pas.

**Figure 8:**
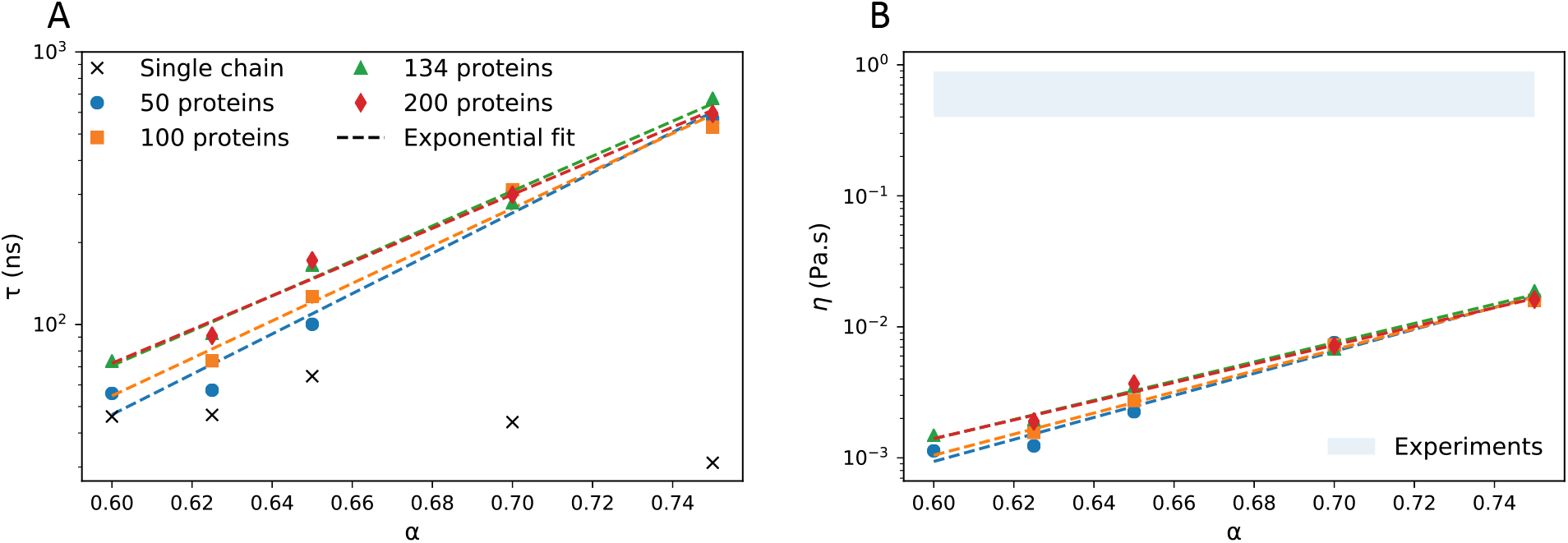
Shear viscosity estimated from the dynamics of the FUS LCD end-to-end distance. (A) Relaxation time *τ* of the end-to-end distance as function of *α* for droplets of different size (filled symbols; as indicated). The crosses show *τ* for an isolated FUS LCD protein free in aqueous solution. (B) Shear viscosity *η* of the droplets as function of α. The blue shaded region indicates the range of experimental estimates reported for FUS LCD.^13,63^ Dashed lines in (A) and (B) are fits to exponential functions in *α*.

## DISCUSSION

### Phase separation

We performed MD simulations of concentrated solutions of FUS LCD using a modified MARTINI coarse-grained model. We scaled the strength of the proteinprotein interactions using the *α* parameter introduced by Stark et al.^40^. For *α* > 0.6, we observed spontaneous phase separation of FUS LCD solutions into a dense phase and a dilute phase. For sufficiently large and dilute systems, the dense regions coalesced into a spherical droplet. In the dilute phase, the FUS LCD chains remained dispersed.

The thermodynamics of phase separation is highly sensitive to the strength of the proteinprotein interactions. In a narrow window of *α* between 0.625 and 0.7, the concentration of the dilute phase varies by about a factor 100 (Figure 4A). In the same window, the dense-phase concentration varies by a factor of three (Figure 4C). As a result, the excess free energy for transferring a FUS LCD chain between the two phases varies by about 15 kJ/mol (Figure 4B) if *α* is changed by 0.1. These sensitivities to seemingly tiny changes in the interaction strength make it clear that force fields have to be rebalanced to describe biopolymer LLPS in a quantitative manner.

The strong dependence on the energy scaling parameter *α* is consistent with the predictions of the van-der-Waals mean-field theory of phase transitions. In Figure 4B, we mapped the van-der-Waals model to the FUS LCD data using a linear relation between the respective attraction parameters a and *α*. The coexistence line of the van-der-Waals model captures the observed phase behavior remarkably well, giving us a rough estimate of the critical value *α_c_* ≈ 0.6. Indeed, below *α* = 0.6, we did not observe phase transition. To the contrary, droplets preformed at higher values of *α* dissolved when *α* was reduced below 0.6, showing that condensation is reversible on the timescale of the MD simulations (Figure S3).

For 0. 65 ≤ *α* ≤ 0. 75, we verified that the initial protein concentration does not influence the protein density in the dense phase, as expected for a phase in thermodynamic equilibrium. Hence, for the simulation of droplets of different sizes, the relevant quantities are the initial number of proteins, the box size, and α. For a given *α* value, the protein concentrations inside droplets of different sizes were remarkably close (Figure 3), again indicating that thermodynamic equilibrium has been established in the simulations.

However, in simulations of systems with initially dispersed proteins at high total concentrations (i.e., close to the dense-phase concentration along the coexistence curve), protein condensates with inverted topologies formed. In these simulations, the dense protein phase was continuous in the periodic system, causing strong finite size effects (Figure 1). Interestingly, for *α* = 0.65, the protein concentration of >400 mg/mL in this percolated dense phase is substantially higher than the ≈300 mg/mL inside the spherical droplets and the periodic slab at the same value of α. Further study of these dense inverted “phases” could be relevant for atomistic simulations of bulk condensates, where box sizes tend to be smaller.

### Optimal interaction strength *α*

In the spirit of the MARTINI model,^39^ we tuned the *α* parameter to reproduce the excess free energy of transferring a FUS LCD chain from the dilute phase into the dense phase according to eq 1. For FUS LCD in aqueous solution at ambient conditions, a value of *α* = 0.65 reproduces the excess transfer free energy, as estimated on the basis of measured densities. Remarkably, for *α* = 0.65 we not only reproduced the excess transfer free energy, which depends on the ratio of the densities in the dense and dilute phases, but also the absolute values of both densities. This served as a first validation test.

The dense-phase concentration increases with *α* and ranges from 150 to 760mg/mL for 0.625 ≤ *α* ≤ 0.8. Experimental studies reported condensed phase concentrations between 120mg/mL^13^ and 285mg/mL.^27^ This second value is close to the concentration we obtained with *α* = 0.65, where our model matched the excess transfer free energy. This result makes us optimistic about the capacity of our model to reproduce also other relevant properties of FUS LCD condensates.

### Effects of droplet curvature

In matching *α* to experiment, we did not take into account surface effects linked to the droplet curvature, which we expect to give rise to a slight increase in the density of the dilute phase. As a rough estimate for *α* = 0.65, we use the Kelvin equation and the ideal gas law to estimate the relative increase in the density of the dilute phase above a droplet of radius *R*,

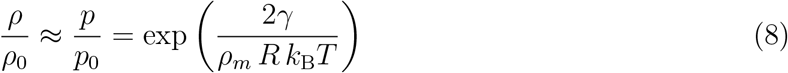

For a density of *ρ_m_* ≈ 0.009 nm^−3^ of FUS LCD in the dense phase, a surface tension of *γ* ≈ 0.05 mN/m, and a radius of *R* ≈ 13 nm, the expected increase in the density of the dilute phase is about 20% and thus comparable to the uncertainties in our estimates of the density of the dilute phase. Conversely, following Powles et al.^65^, we solved eq 8 for the surface tension *γ*, with *ρ* the density of the dilute phase above the droplets, *ρ*_0_ the density of the dilute phase above the slab (Figure S11), *ρ_m_* the density of the droplets, and R the radius of the droplets. For *α* = 0.65, we obtained surface tensions of 0.13 to 0.27mN/m, about two to six times higher than our estimates from droplet shape fluctuations. In light of the simplifications in eq 8, where we assumed the dilute phase to be ideal, and the uncertainties from finite-size effects and slow convergence of the dilute-phase concentration, we find this semi-quantitative agreement in the calculated surface tensions to be reassuring. We refer to ref 17 for a detailed discussion of curvature effects.

### Slow relaxation at high *α*

We calculated the protein mass density inside the droplets and monitored for possible inhomogeneities. For 0.65 ≤ *α* ≤ 0.75, the dense phase appeared homogeneous, consistent with a liquid-like state of these droplets. By contrast, for *α* ≥ 0.8, significant inhomogeneities emerged. At these high values of *α*, FUS LCD condensed and then got trapped in structures resembling amorphous aggregates subject to slow relaxation on the MD timescale. Extrapolating to such high values of *α*, the end-to-end distance relaxation occurs on timescales of several microseconds (Figure 8A). Such slow chain reconfiguration and the associated high viscosity of the dense phase prevent the relaxation on an MD timescale for *α* ≥ 0.8. Patel et al.^9^ reported that physiologically competent FUS droplets have to be liquid, but that aging can lead to aggregation and transition to a solid-like state. By performing long MD simulations of FUS LCD condensates for *α* ≥ 0.8 (or gradually increasing *α*), it might be possible to mimic aspects of droplet aging and monitor changes in the apparent materials properties.

### Hydration of droplet interior

Thanks to the explicit treatment of the solvent by the MARTINI model, we could quantify the droplet hydration levels. The computed water mass fraction profiles inside the droplets showed that water accounts for ≈ 70% of the droplet mass at *α* = 0.65 (Figures 3B and 5). As *α* is increased, the water content of the droplets decreases, reaching ≈50% for *α* = 0.75. In vivo, water is critical to maintain the fluidity of the droplet, the diffusivity of molecules within, and their biochemical reactivity.^66^ Murthy et al.^27^ report a water content of ≈ 65% by volume for FUS LCD droplets. The close correspondence with the calculated water content again suggests *α* = 0.65 as a reasonable rescaling parameter.

### Surface tension

We characterized the droplet surface tension in terms of droplet shape fluctuations and interfacial widths. The three estimates obtained in this way for a given value of *α* are in excellent correspondence (Figure 6). We found that the surface tension increases strongly with *α* over a narrow range. Indeed, as the critical value *α_c_* ≈ 0.6 is approached, the surface tension drops to zero, consistent with theoretical expectations (Figure 6). As *α* is increased, strengthened protein-protein interactions increase the cohesive forces and, in turn, the surface tension.

For 0.625 ≤ *α* ≤ 0.75, we obtained surface tensions ranging from 0.015 to 0.38 mN/m. Taylor et al.^67^ reported surface tensions of 0.004mN/m and 0.68mN/m for the IDPs Whi3 and LAF1, in the range of our estimates for FUS LCD.

For *α* = 0.65, where the experimental densities of the dilute and dense phases of FUS LCD are reproduced, the surface tension is about 0.05 mN/m. For reference, the water-vapor surface tension at ambient conditions is about 1500 times larger. Even the liquid-vapor surface tension of a LJ fluid at a corresponding state is more than 100 times larger. At a reduced temperature of 0.85, where the excess transfer free energy of the LJ fluid matches that of our FUS LCD model at *α* = 0.65, a reduced surface tension of about 0.837 has been reported,^68^ corresponding to γ = 6.7 mN/m. Here we used effective LJ parameters *σ* = 0.78 nm and *ϵ* = 2.9kJ/mol for which the densities at coexistence^68^ match those of FUS LCD (Figure 3). The wide gap between the surface tensions of FUS LCD and the corresponding LJ fluid indicates that for interfacial properties the polymeric nature of the droplets and the immersion in an aqueous solvent cannot be neglected.

In an effort to rationalize the low surface tensions obtained here and reported from experiment, we note that the increase in the surface tension goes along with a decrease in the droplet water content (Figure S12). For large *α*, we expect the surface tension to saturate at values for compact and dry protein globules (estimated^69^ at ≈50mN/m). The extrapolation of the exponential fits of *γ* to zero water content gives a surface tension of 10 to 20mN/m for a compact protein (Figure S12). So, at least roughly, we are indeed interpolating between a compact phase (water content zero) and a dissolved phase (*γ* = 0; water content 100%). Note that the exponential dependence on *α* has to break down as the dissolved phase and the critical *α* are approached, where *γ* = 0.

### Shear viscosity

We also estimated the shear viscosity of the FUS LCD droplets. We found the viscosity to increase exponentially with *α* from 0.001 to 0.02Pas for 0.6 < *α* < 0.8. For *α* = 0.65, we found *η* ≈ 0.004Pas. For reference, Murakami et al.^63^ reported a viscosity of 0.4Pas, and Burke et al.^13^ reported a viscosity of ≈0.9Pas. These values are larger than our estimate by about a factor of one hundred. By performing further comparisons for other proteins undergoing LLPS, it will be interesting to see if the substantially lower viscosity of the MD model is inherent to coarse-graining the potential energy surface, which tends to smoothen the strong distance and orientation dependence of atomic interactions.

### Transferability

We emphasize that the optimal value of the scaling parameter determined here, *α* = 0.65, applies only to FUS LCD under the thermodynamic conditions of the present study. For other phase separating systems, the optimal values of *α* may be different. Nevertheless, one may hope that the value of *α* = 0.65 obtained here for FUS LCD serves as a rough estimate also for other proteins, which would give the rebalancing procedure a degree of transferability. Conversely, explorations of the connection between the optimal values of the scaling parameter *α* and the nature of the proteins should give insight into the molecular properties that favor phase separation.

### Interaction rebalancing in LLPS modeling

Our rebalancing approach is general and applicable to different types of coarse-grained simulation models of phase separation. There has been much progress in devising coarse-grained^70–72^ and implicit solvent models^73^ of disordered proteins and we envisage that our general approach will make it possible to leverage these developments in coarse-grained modeling of disordered biomolecules for simulations of LLPS and biomolecular condensates. We note that our tuning of the relative strengths of protein-protein and protein-solvent interactions is similar to corrections implemented for highly optimized atomistic protein force fields.^35–37,74^ Further fine tuning may be required for quantitative atomistic simulations of LLPS.

The strong dependence of the droplet biophysical properties on *α* underlines how molecular interactions have an impact on physical properties on larger length scales, as it has been stressed before. ^75^ In experiment, salt or condensing factors are added to modulate the interaction strength. At least at a qualitative level, changes of *α* in the simulations may thus mimic these changes in solvent conditions of the experiments. As we start to use MD simulations to study phenomena of increased complexity and on a larger scale, rebalancing may thus be both a necessity and an opportunity: a necessity to avoid the large errors resulting from systematic imbalances, and an opportunity to probe the phase behavior over a wide range with only minimal modifications to the system.

## ASSOCIATED CONTENT

### Supporting Information

Detailed derivation of the relations between surface tension and droplet shape fluctuations, and between surface tension and droplet interfacial width; amino acid sequence of FUS LCD; table of simulations; droplet density inhomogeneities; protein cluster formation; reversibility of phase separation; radial protein density profiles; droplet shape fluctuations; cumulative distributions of the amplitudes of shape fluctuations; autocorrelation functions of end-to-end distance of FUS LCD chains; average end-to-end distance autocorrelation functions; distributions of end-to-end distances; determination of diffusion coefficient for end-to-end distance; and connection between surface tension and hydration.

Supporting Movies S1 and S2, showing phase separation of FUS LCD at *α*=0.7 and water inside phase-separated FUS LCD droplet.

## Acknowledgement

We acknowledge financial support from the German Research Foundation (CRC 902: Molecular Principles of RNA Based Regulation), the Human Frontier Science Program RGP0026/2017 (S.v.B. and G.H.), and the Max Planck Society. The authors thank Drs. Mateusz Sikora and Jurgen Köfinger as well as Lisa M. Pietrek for insightful and helpful discussions.

## Supporting Information

### Supporting Text

#### Surface tension from droplet shape fluctuations

Following Henderson and Lekner^58^ we assume that the energetics underlying the thermal fluctuations in the droplet shape is dominated by the changes in surface area. The potential energy *U* of such surface-shape fluctuations is dominated by the surface tension *γ*:

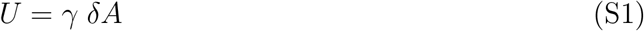

where *δA* is a small change in surface area relative to the sphere. The fluctuations in the shape of the droplet at lowest order in a spherical harmonics expansion give us two independent estimates of the surface tension, The surface of the droplet can be described as a sum of spherical harmonics 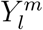:

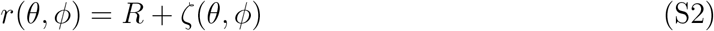

with *r* the radius as a function of the polar and azimuthal angles *θ* and *φ*, and

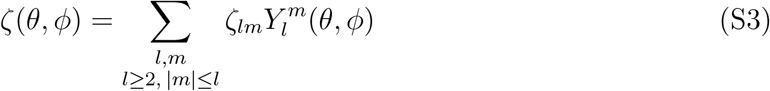

*R* is the average radius and *ζ_lm_* is the coefficient of the mode (*l, m*). For small amplitudes, the potential energy *U* then becomes:

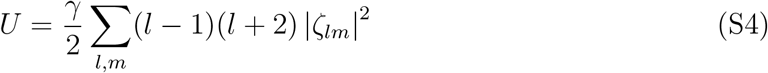

For small perturbations (|*ζ*|^2^ ≪ *R*), each mode (*l, m*) is thus effectively harmonic and independent. As a consequence, the equipartition theorem gives a relation between the surface tension and the mean squared amplitudes of the shape fluctuations:

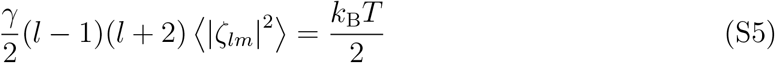

Each mode (*l,m*) gives an independent estimate for the surface tension. Focussing on the low-frequency modes, we approximate the instantaneous droplet shape as an ellipsoid with axes *a, b* and *c*. If this ellipsoid is aligned with the Cartesian coordinate system, its spherical coordinate representation is:

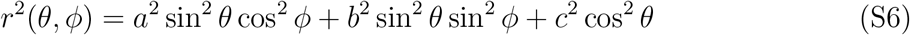

We expand *r*(*θ, ϕ*) to second order in *a* = *R* + *δa, b* = *R* + *δb, c* = *R* + *δc*. By projecting onto spherical harmonics, we find:

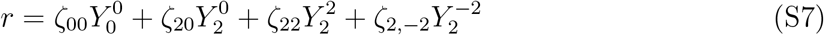

where 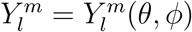 and *r* = *r*(*θ, ϕ*). The expansion coefficients are:

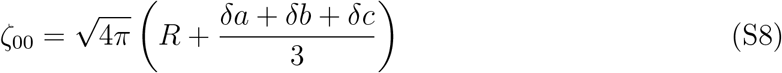

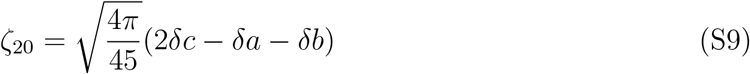

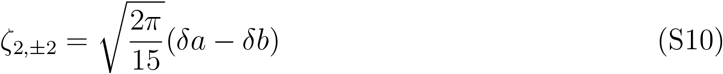

The condition of constant volume requires that:

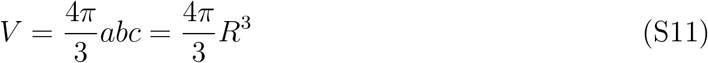

To lowest order, we have

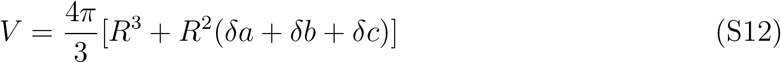

and thus

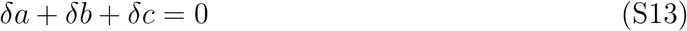

By imposing this condition on the amplitudes of the modes, we find

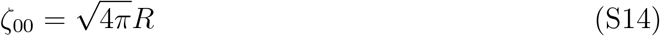

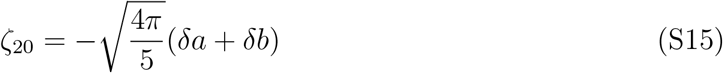

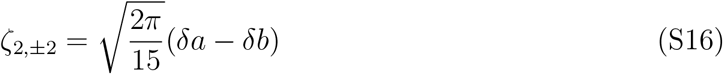

We thus arrive at an expression for the potential energy associated with surface area changes *U* = *γ δA* to second order in a spherical harmonics expansion:

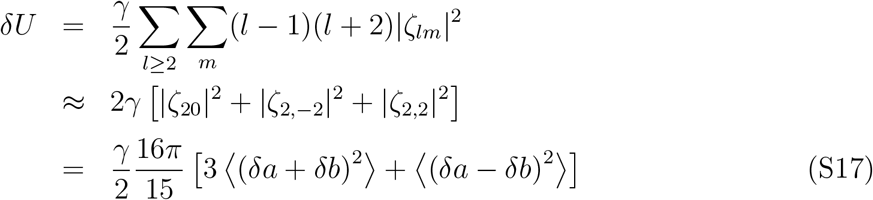

This expression for the potential energy combined with the equipartition theorem for the two independent modes, *δa* + *δb* and *δa* – *δb*, gives us the two independent expressions for the surface tension, eqs 3 and 4.

##### Alternative derivation of relation between surface tension and shape fluctuations

The preceding expression for the surface tension can also be derived directly from eq S1. We again describe the instantaneous droplet shape by an ellipsoid with axes *a, b*, and *c*. We expand the surface area of the ellipsoid to second order in *δa, δb* and *δc*. In terms of spherical polar angles *θ* and *ϕ*, the first fundamental form defining the surface area element can be written as

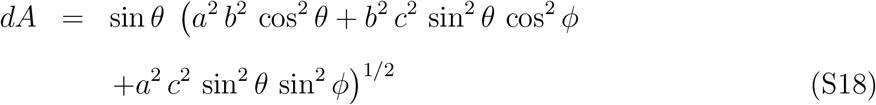

The area of the ellipsoid then becomes

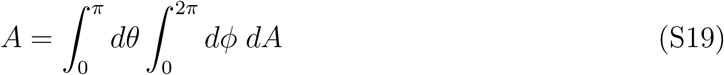

In our statistical mechanical model of droplet shape fluctuations, we (1) impose droplet volume conservation by setting *c* = *R*^3^/(*ab*), (2) introduce new variables *a* = *R* + (*δu* + *δv*)/2 and *b* = *R* + (*δu* – *δv*)/2, (3) expand the area element *dA* to second order in *δu* and *δv*, and (4) integrate over the polar angles to obtain an expression for the area of the ellipsoid to second order in *δu* and *δv*. In this way, we arrive at

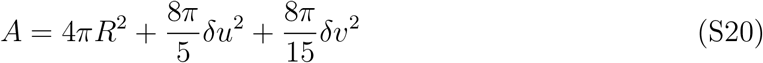

Substituting the area change *δA* = *A* – 4*πR*^2^ into eq S1 for the corresponding potential energy, we find that the fluctuations in *δu* = *δa* + *δb* and *δv* = *δa* – *δb* are thus harmonic and uncoupled. The equipartition theorem combined with the surface energy eq S1 then gives us the expressions for the surface tension in eqs 3 and 4.

#### Droplet shape

We described the droplet shape in terms of a general ellipsoid with axis lengths *a, b, c*. We estimated *a, b* and *c* from principal component analysis (PCA) of the mass distribution. Let ***r***_CMS_ be the center of mass of the atoms *i* with positions ***r**_i_* and masses *m_i_* in the droplet (excluding water, ions and other small molecules in the solvent):

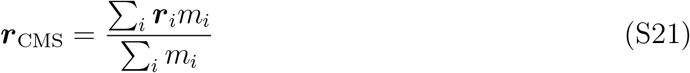

For simplicity, we gave all protein beads in MARTINI an equal mass weight *m_i_* = 1. We then calculated a 3×3 mass-weighted covariance matrix of the Cartesian positions 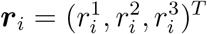 with elements

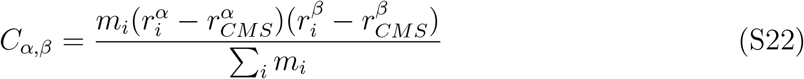

The eigenvalues *λ*_1_, *λ*_2_, and *λ*_3_ of the matrix ***C*** are proportional to the squared ellipsoidal axes: *λ*_1_ = *νa*^2^, *λ*_2_ = *νb*^2^, *λ*_3_ = *νc*^2^. By imposing the condition of constant volume *R*^3^ = *abc*, we eliminate *ν*:

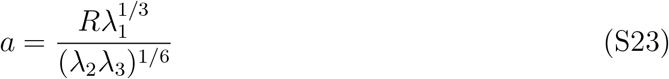

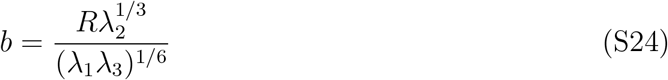

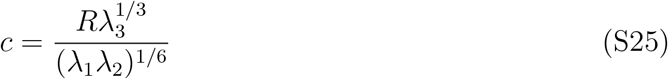

where *R* is the average radius of the droplet, obtained from a sigmoidal fit of the density profiles (eq 2). If the instantaneous droplet shape has ellipsoidal axes *α*_1_(*t*), *α*_2_(*t*), *α*_3_(*t*) at time *t* along an MD trajectory, the instantaneous axis fluctuations are then *δa_i_*(*t*) = *a_i_*(*t*) – *R*. From the deviations of the instantaneous axes from the mean droplet radius averaged over the three independent combinations of axes, we estimate the variances

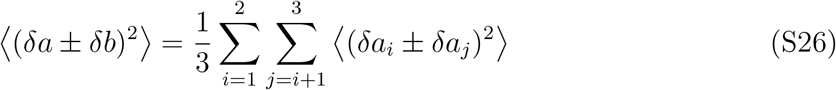

Substituting these averages into eqs 3 and 4, we arrive at the following two independent estimates of the surface tension

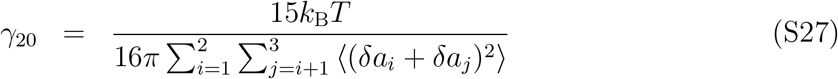

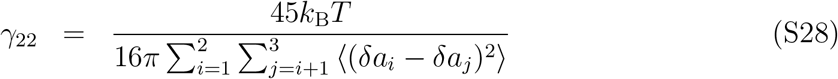

We expect that the surface-tension estimates are consistent, *γ* = *γ*_20_ = *γ*_22_.

### Supporting Tables

**Table S1:** Amino-acid sequence of the 163-residue FUS low complexity domain.

|  |  |  |  |  |
| --- | --- | --- | --- | --- |
| 1-MASNDYTQQA | TQSYGAYPTQ | PGQGYSQQSS | QPYGQQSYSG | YSQSTDTSGY-50 |
| 51-GQSSYSSYGQ | SQNTGYGTQS | TPQGYGSTGG | YGSSQSSQSS | YGQQSSYPGY-100 |
| 101-GQQPAPSSTS | GSYGSSSQSS | SYGQPQSGSY | SQQPSYGGQQ | QSYGQQQSYN-150 |
| 151-PPQGYGQQNQ | YNS-163 |  |  |  |

**Table S2:** Simulation characteristics. Listed are the starting configuration, the width of the cubic box, the number of FUS LCD chains, their overall concentration, and the *α* interaction rescaling parameters. Slab box size “*” corresponds to 90 × 20 × 20 nm^3^. A prime in the *α* column indicates a run length of 12 μs instead of 24 μs.

| Starting configuration | Box width (nm) | Number of proteins | Concentration (mg/mL) | Time ($\mu\text{s}$) | $\alpha$ |
| --- | --- | --- | --- | --- | --- |
| Homogeneous | 40 | 134 | 60 | 12 | 0.1, 0.2, 0.3, 0.5, 0.55, 0.6, 0.65, 0.7 |
|  |  | 336 | 150 | 12 | 0.1, 0.2, 0.3, 0.5, 0.55, 0.6, 0.65, 0.7, 0.75, 1 |
|  |  | 672 | 300 | 12 | 0.1, 0.2, 0.3, 0.5, 0.55, 0.6, 0.65, 0.7, 0.75, 0.8, 1 |
| Preformed droplet | 40 | 134 | 60 | 12 | 0.5, 0.55, 0.65, 0.7, 0.75, 0.8, 0.85 |
|  | 50 | 50 | 11 | 24 | 0.6', 0.625', 0.65', 0.7, 0.75' |
|  |  | 100 | 23 | 24 | 0.625, 0.65', 0.7, 0.75' |
|  |  | 134 | 31 | 24 | 0.6', 0.625, 0.65, 0.7, 0.75' |
|  | 60 | 200 | 26 | 24 | 0.625, 0.65, 0.7, 0.75 |
| Single FUS chain | 20 | 1 | 3.6 | $6 \times 5$ | 0.6, 0.625, 0.65, 0.7, 0.75 |
| Slab | * | 100 | 79 | 36 | 0.6, 0.625, 0.65, 0.7 |

**Table S3:** Trajectory segments used for the surface tension calculation (in μs) for simulations with different numbers *N* of FUS LCD proteins and different scaling factors *α*. In these segments, the droplet had relaxed and, at *α* = 0.625, was clearly discernible.

| $\alpha$ | Shape fluctuations | | | | Interfacial width | | | |
| --- | --- | --- | --- | --- | --- | --- | --- | --- |
| | $N = 50$ | 100 | 134 | 200 | $N = 50$ | 100 | 134 | 200 |
| 0.625 | – | – | – | – | 6-12 | 6-12, 22-24 | 6-24 | 6-24 |
| 0.65 | 10-12 | 10-12 | 6-24 | 6-24 | 6-12 | 10.5-12 | 16-24 | 18-24 |
| 0.7 | 16-24 | 16-24 | 15-24 | 6-24 | 16-24 | 15-24 | 6-24 | 6-24 |
| 0.75 | 4-12 | 7-12 | 6-12 | 11-12 | 8.7-12 | 10-12 | 8.7-12 | 6-12 |

### Supporting movies

**Movie S1:** Phase separation of FUS at *α* = 0.7 leading to the formation of a spherical droplet starting from a homogeneous solution of *N* = 134 proteins in a 40 × 40 × 40 nm^3^ box. The system adopts different structures, being first a “tube” spanning the box that then ruptures to form a sphere. Shown are the FUS LCD chains in different colors, with water and ions omitted for clarity. The length of the shown trajectory is 12 μs.

**Movie S2:** Water inside phase-separated FUS droplet for *α* = 0.7 with *N* = 134 proteins. In this MD simulation snapshot, FUS LCD proteins are shown in red, and water beads in light blue. A clip plane for the protein is moved through the droplet, leaving behind the exposed water beads to reveal the hydration gradually, ending with a “ droplet of water”.

### Supporting Figures

**Figure S1:**
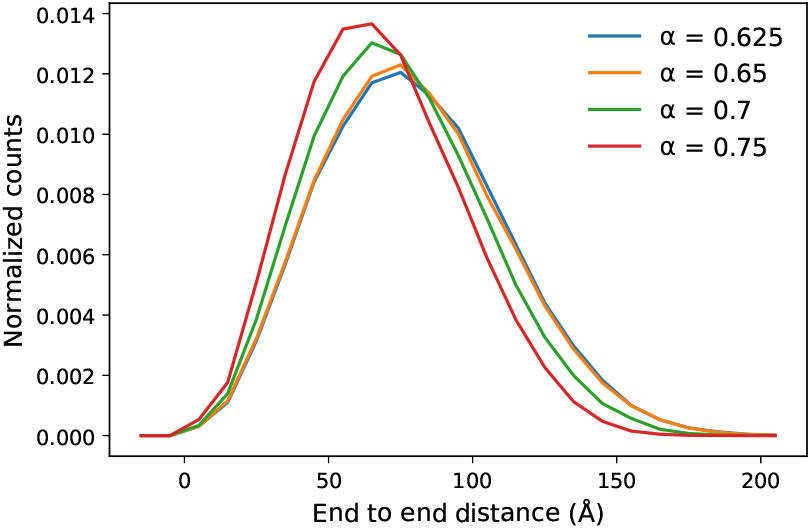
Distribution of end-to-end distances at various values of *α* in MD simulations with *N* = 134 proteins.

**Figure S2:**
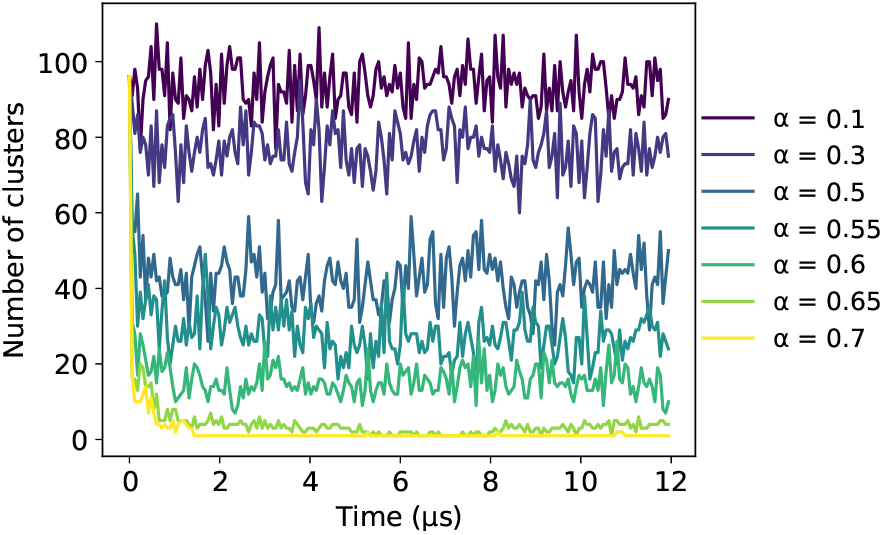
Cluster formation. The number of FUS LCD clusters is plotted as function of time for different *α*. The simulations started from a homogeneous solution of 134 proteins in a 40 × 40 × 40 nm^3^ box, equilibrated with *α* = 0. During the first 2 μs, following the increase of *α* from zero to the desired value at time zero, larger clusters formed and, as a result, the overall number of clusters decreased.

**Figure S3:**
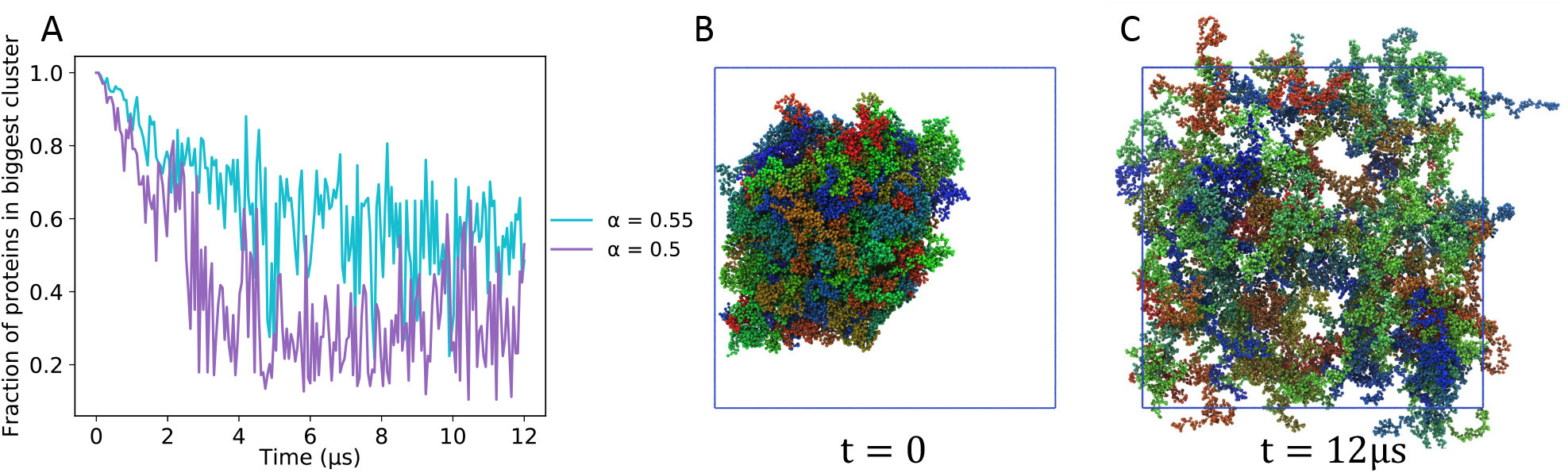
Reversibility of the FUS LCD phase separation. (A) Fraction of proteins in the largest cluster as a function of time for *α* = 0.5 (magenta) and *α* = 0.55 (light blue). The MD simulations started with a droplet preformed at *α* = 0.7 with *N* = 134 proteins. This droplet dissolved after *α* was lowered to 0.55 or 0.5. (B) Starting configuration with a preformed droplet. (C) Final configuration at *α* = 0.5 with dispersed FUS LCD.

**Figure S4:**
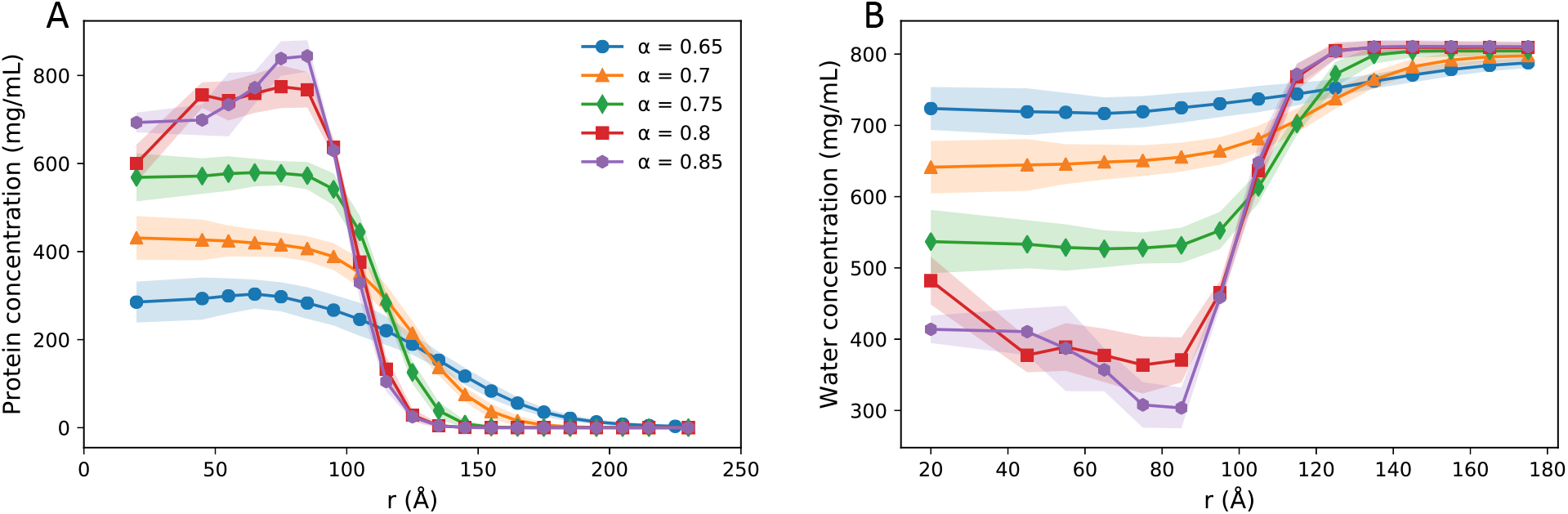
Protein concentration inhomogeneities in FUS LCD droplets for *α* > 0.75. (A) Radial protein density profiles for *α* between 0.65 and 0.85. (B) Corresponding water density profiles.

**Figure S5:**
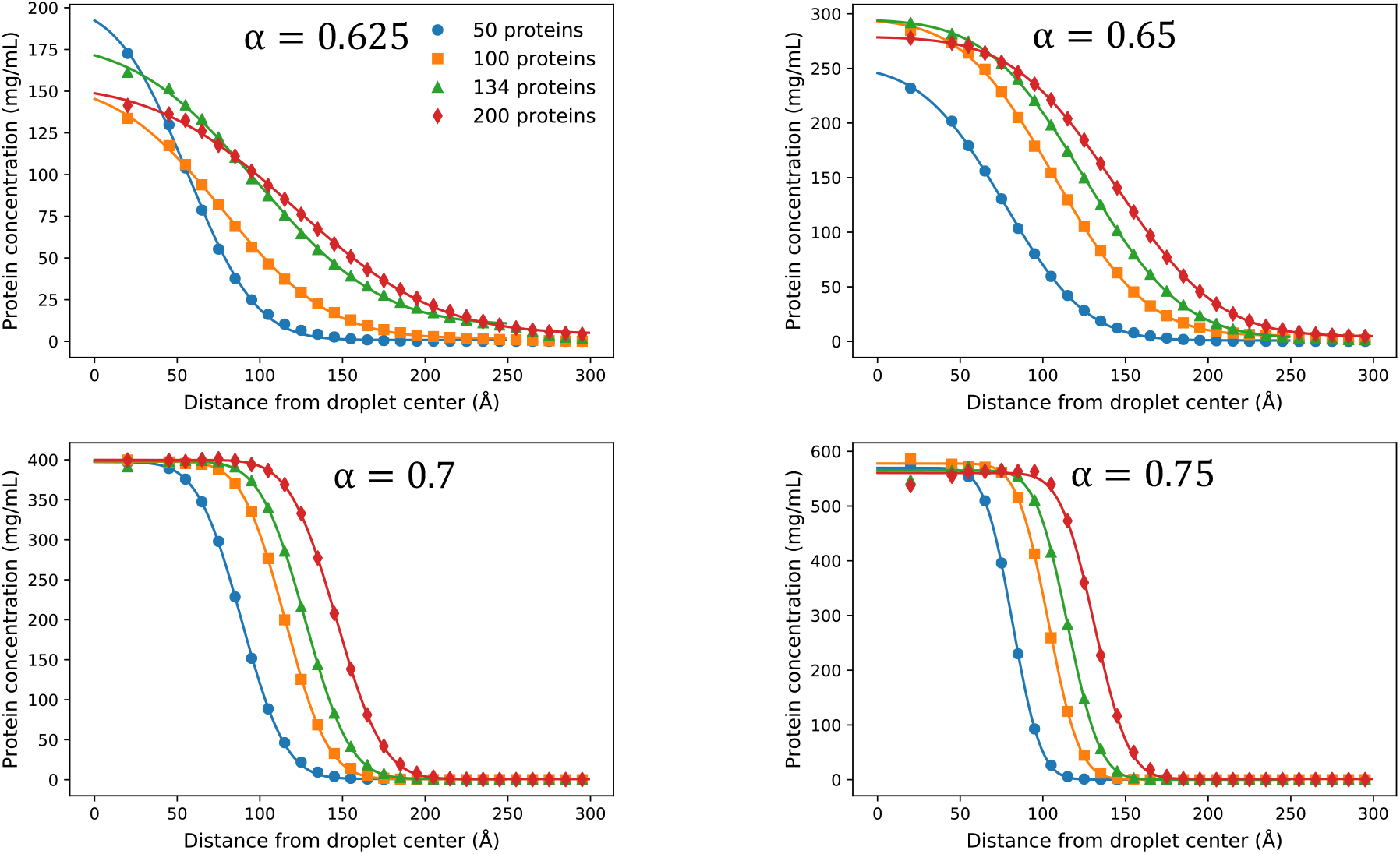
Radial protein concentration profiles as function of distance from droplet center (symbols: simulation results; lines: fit to error function density profile). Results are shown for MD simulations with different total numbers *N* of proteins in a box of volume 50 × 50 × 50 nm^3^ for *N* = 50, 100, 134 proteins and 60 × 60 × 60 nm^3^ for *N* = 200 proteins, and for different values of *α*.

**Figure S6:**
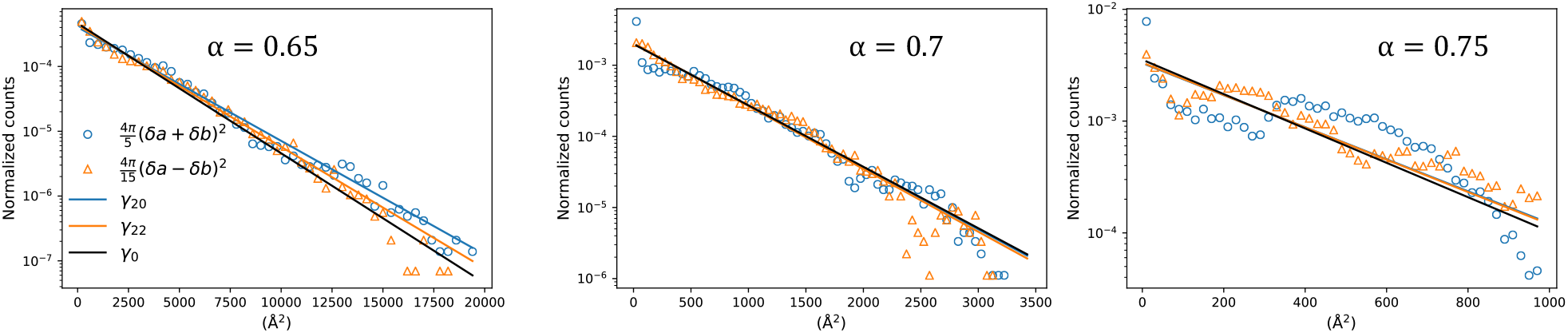
Droplet shape fluctuations. Shown are the normalized histograms of the scaled ellipsoidal mode amplitudes, |*ζ*_20_|^2^ and |*ζ*_22_|^2^ + |*ζ*_2,-2_|^2^, for a droplet made of 134 proteins at different values of *α*. The lines are predictions from the capillary wave models, using the different estimates of the surface tension as input, as indicated. Deviations at high values of *α* (right) are likely the result of slow shape relaxations of the viscous drops on the MD timescale.

**Figure S7:**
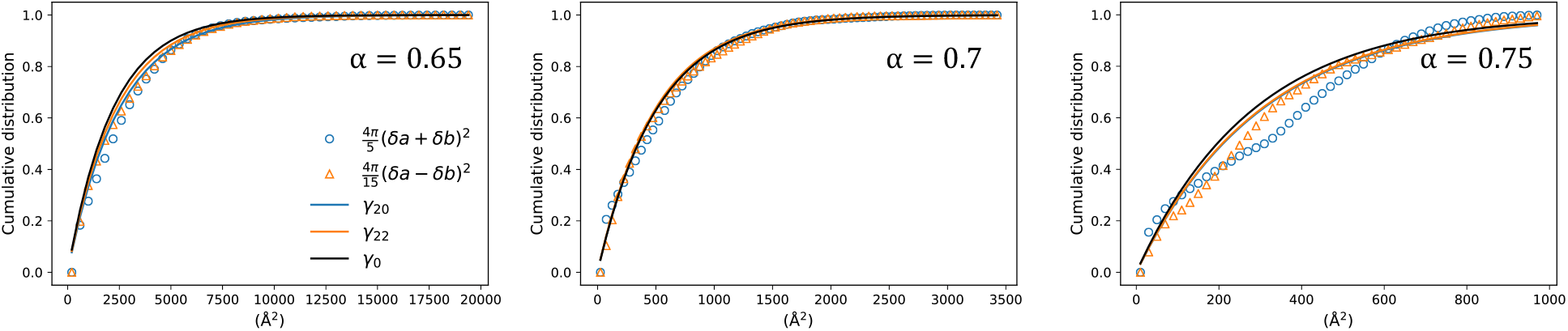
Cumulative distributions of the scaled ellipsoidal modes amplitude |*ζ*_20_|^2^ and |*ζ*_22_|^2^ + |*ζ*_2,-2_|^2^, for a droplet made of 134 proteins at different *α*. The lines are predictions from the capillary wave models. See Figure S6 for the corresponding histograms.

**Figure S8:**
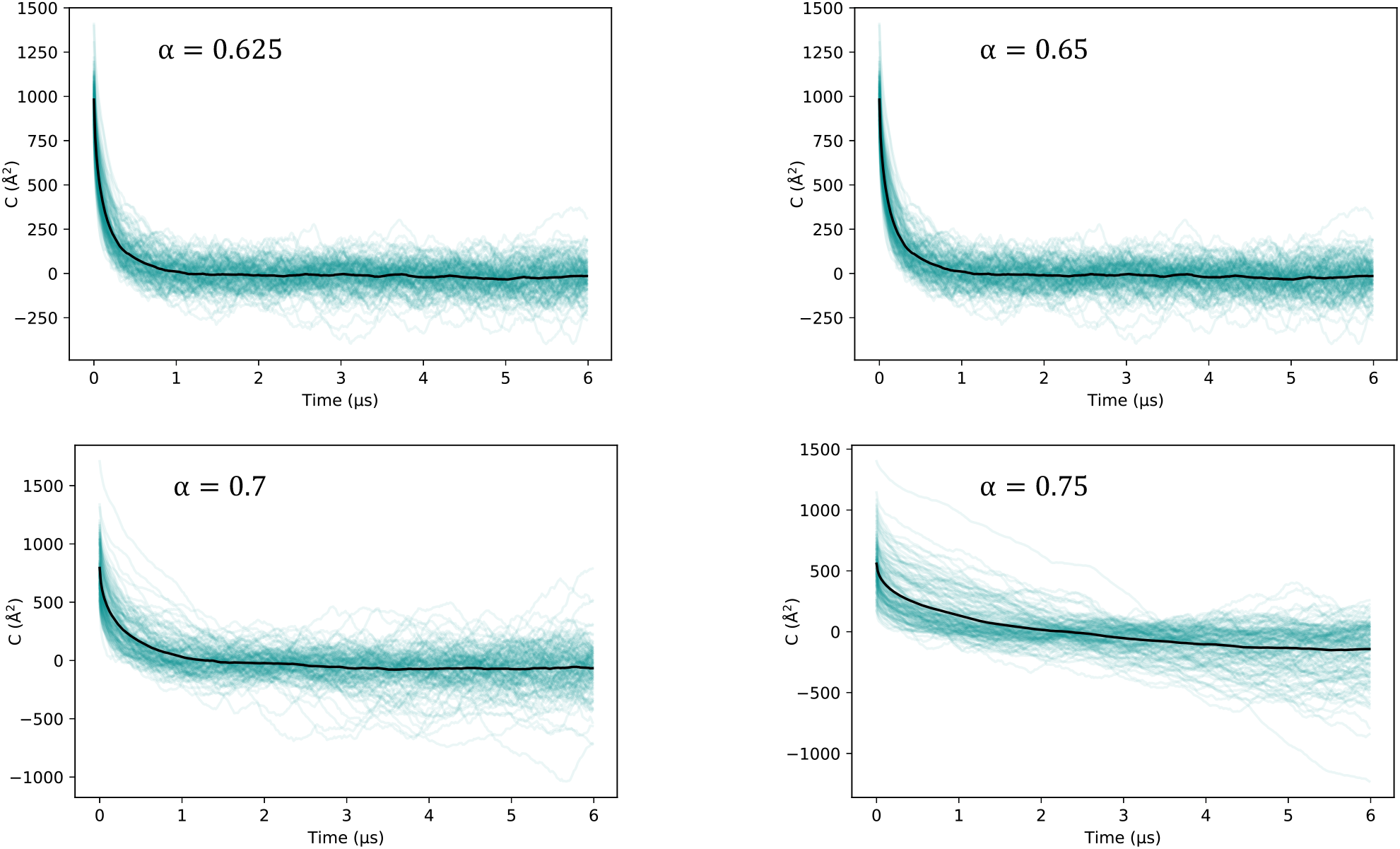
Autocorrelation functions of the FUS LCD end-to-end distance at different values of α. Thin turquoise lines are the results for individual FUS LCD chains. The average of the autocorrelation functions across the ensemble of 134 protein chains is shown in black.

**Figure S9:**
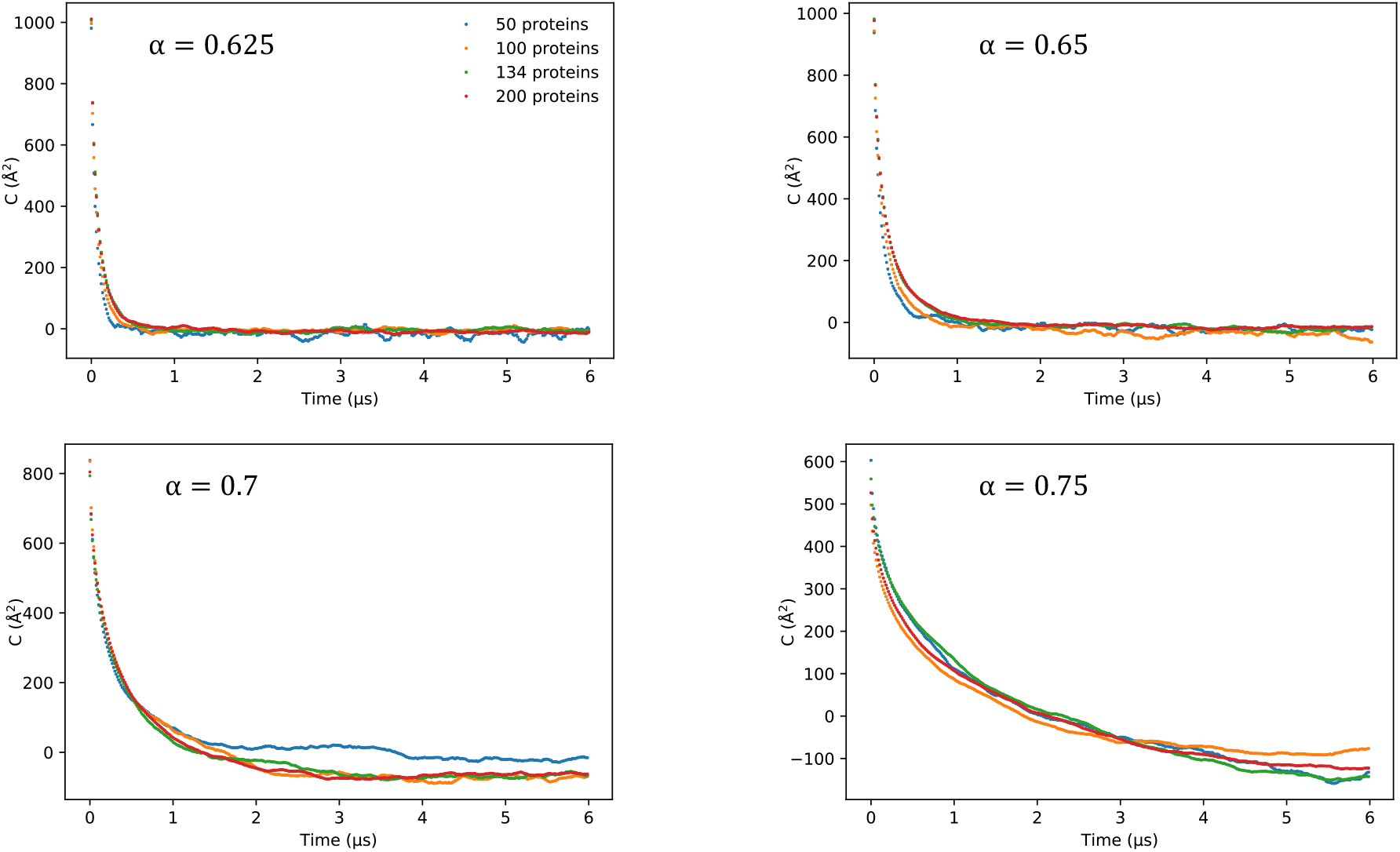
Average of the FUS LCD end-to-end distance autocorrelation function across the ensemble of chains at various values of *α* and for different numbers of proteins.

**Figure S10:**
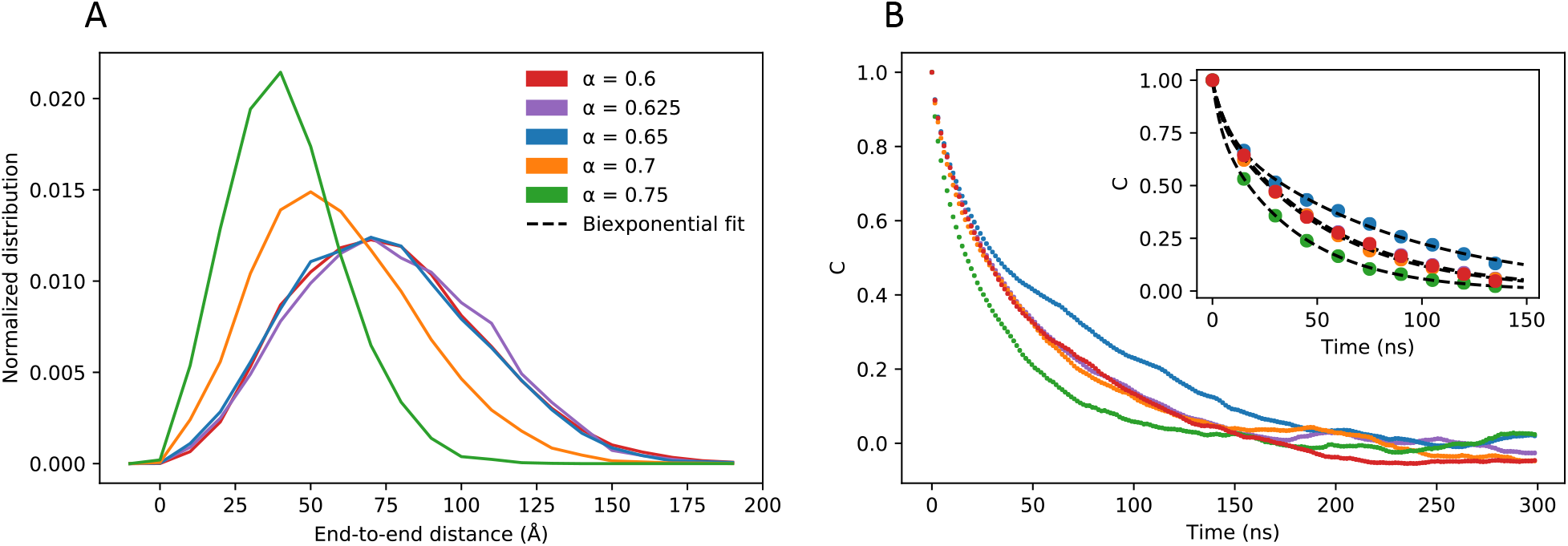
Determination of effective diffusion coefficient for end-to-end distance of isolated FUS LCD chains. (A) Normalized distributions of the end-to-end distances of an isolated chain free in solution at different values of α. Note that chain compaction sets in for *α* ≥ 0.7. (B) Autocorrelation functions of the end-to-end distances of isolated chains, averaged over five independent runs of 6 μs each. The inset shows the biexponential fits from which the relaxation times were computed (crosses in Figure 8).

**Figure S11:**
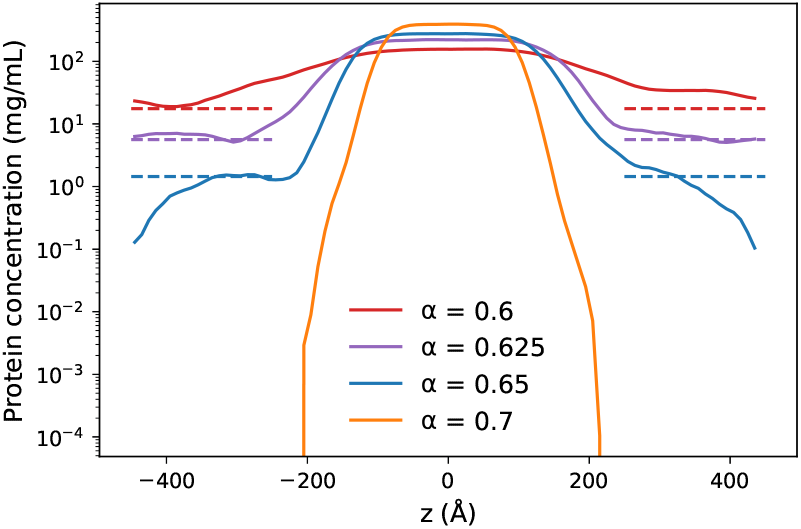
Slab concentration profiles. The dashed lines show the dilute-phase concentrations obtained from clustering. Results are shown for MD simulations with 100 proteins in a box of volume 20 × 20 × 90 nm^3^ and for different values of *α*, as indicated.

**Figure S12:**
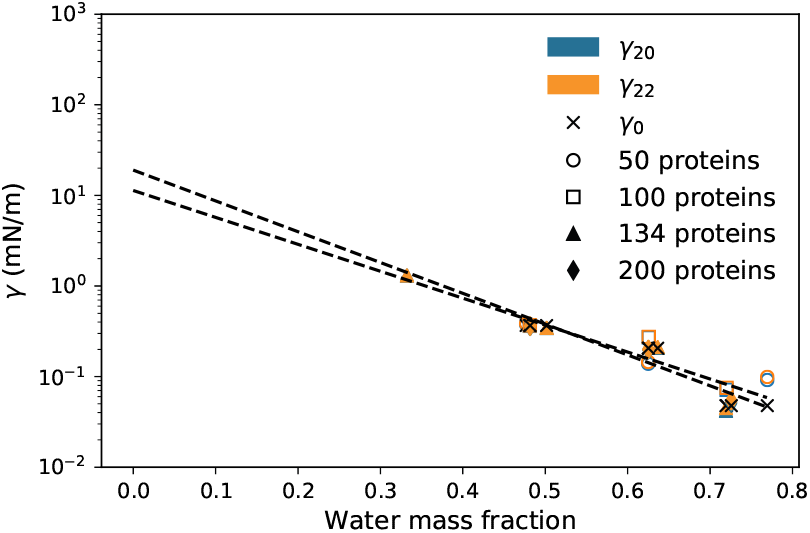
Connection between surface tension and hydration. Results are shown for *α* = 0.8, 0.75, 0.7 and 0.65, which appear as distinct groups of points from left to right. The black dashed lines are exponential fits to *γ*_20_ and *γ*_22_, and to *γ*_0_, in an effort to extrapolate the surface tension to zero water content (i.e., a dry droplet).

